# Plant pathogen infection risk tracks global crop yields under climate change

**DOI:** 10.1101/2020.04.28.066233

**Authors:** Thomas M. Chaloner, Sarah J. Gurr, Daniel P. Bebber

## Abstract

Global food security is strongly determined by crop production. Climate change-induced losses to production can occur directly, or indirectly, including via the distributions and impacts of plant pathogens. However, the likely changes in pathogen pressure in relation to global crop production are poorly understood. Here we show that temperature-dependent infection risk, r(*T*), for 80 fungal and oomycete crop pathogens will track projected yield changes in 12 crops over the 21st Century. For most crops, both yields and r(*T*) are likely to increase at high latitudes. In contrast, while the tropics will see little or no productivity gains, r(*T*) is also likely to decline. In addition, the USA, Europe and China may experience major changes in pathogen assemblages. The benefits of yield gains may therefore be tempered by the increased burden of crop protection due to increased and unfamiliar pathogens.

## Main text

Plant pests and pathogens exert a major burden on crop production around the world ^1^. The burden can be measured directly in yield losses or indirectly in the social, environmental and economic costs of control ^1^. Like all species, crop pests and pathogens have particular tolerances to, or requirements for, particular environmental conditions ^2^. These tolerances define their ecological niche, which determines the geographical regions and periods of the year that allow pests and pathogens to proliferate and attack crops ^2^. As climate changes, suitable conditions for pest outbreaks shift in time and space, altering the threats that farmers face and the management regimes required for their control ^3^. Modelling the pattern and process of future changes in pest and pathogen burdens is therefore a key component in maintaining future food security ^4^.

Latitudinal range shifts of pests and pathogens are expected as the planet warms and populations track their preferred temperature zones ^3^. Spatial movements in geographical distributions and temporal shifts in phenologies of wild populations are among the clearest signs of anthropogenic global warming ^5^. Though distribution data for crop pests and pathogens are noisy and incomplete ^4^, similar changes have been detected for hundreds a species of pests and pathogens over recent decades ^6^. Increasing burdens of insect pests at high latitudes, and decreasing burdens at low latitudes, have been projected using ecological niche models (ENM) ^7^. ENMs attempt to reconstruct the environmental tolerances of species from contemporary climates within the observed species range using statistical models ^8^. Alternatively, species’ responses to microclimate can be directly measured, and these responses incorporated into physiologically-based models of species performance ^9^. Such mechanistic models are commonly used to project future crop yields ^10^, and models have also been developed for some plant diseases ^11,12^. However, we know little about how plant disease pressure is likely to change in future, nor how these changes will relate to crop yield responses to climate change.

Infection and disease are determined by complex and species-specific interactions between various biotic and abiotic factors^1^. Temperature is a major determinant of disease risk ^2,13^ and global distributions of plant pathogens have shifted in line with historical global warming ^6^. Here, we analyse temperature response functions for host infection for a suite of fungal and oomycete plant pathogens. We model the likely global shifts in temperature-dependent infection risk for the 21^st^ Century and compare climate-driven changes in this risk with projected changes in crop yields.

### Projected crop yield changes

We compared current (2011-2030 mean) and future (2061-2080 mean) yields projections from three crop models (LPJmL, GEPIC, PEPIC) employing four GCMs (GFDL-ESM2M, HADGEM2-ES, IPSL-CM5A-LR, MIROC5) under the RCP6.0 representative concentration pathway. Carbon dioxide fertilization effects were included, and we compared projections with and without irrigation. Crop models do not explicitly consider the impacts of pests, pathogens and weeds on production. The major commodity crops of maize, wheat, soybean and rice are considered in all three crop models.

Crop models project greater yield increases at higher latitudes, with smaller increases or yield declines at low latitudes^14,15^ (Fig. S1-S4). Under the no irrigation scenario, GEPIC/PEPIC project substantial maize yield declines in Central and Latin America except for Argentina, and across Africa and northern Australia. LPJmL projects no such yield declines. Wheat yields also increase at high latitudes in all three crop models, with smaller increases at low latitudes in LPJmL and declines in GEPIC/PEPIC. North America and parts of Eurasia show the largest wheat yield increases, while GEPIC projects large declines in yield across the tropics. A similar latitudinal trend is projected for soybean but with little decline in the tropics. Soybean yield increases are projected across Eurasia in all models, and also Argentina and South Africa in GEPIC/PEPIC. The latitudinal gradient is less pronounced for rice, with the MIROC5 climate model suggesting a large increase in yield in the Southern hemisphere.

Eight further temperate and tropical annual crops are considered in LPJmL. In the unirrigated scenario, cassava yields increase under all four GCMs within 40 ° of latitude, driven by large increases in India. However, all four GCMS suggest a smaller increase within 10 °N, caused by a yield decline in northern Brazil. Peanut, pea, rapeseed, sugarbeet, and sunflower show increases at all latitudes, with the largest increases at higher latitudes. Millet also shows increases at high latitudes, but yield declines at low latitudes. There are no consistent differences among the four GCMs for any of the crops. Results for sugarcane are more variable. Mean yield change projections suggest declines in Brazil and other Latin American countries, and in Southeast Asia, but an increase in the USA and in East Africa. Previous analyses based on the more extreme RCP 8.5 scenario similar yield increases with latitude latitudes, but more severe declines for some crops at low latitudes ^15^.

Total projected crop production change is difficult to estimate because the spatial distributions of planted areas are impossible to predict, due to the influence of socioeconomic and cultural factors on planting choice. However, if production is calculated from projected yield changes on an estimate of current crop production, increases in production are expected for many crops (Fig. S5). Global wheat, cassava, rapeseed and sunflower production are predicted to increase by all models. LPJmL, and two climate models driving GEPIC/PEPIC, predict increases for rice. All models except HADGEM2-ES predict global soybean production increases (see Methods for analysis of soybean production). None of the crop models unequivocally project declines in production for any crop. In summary, crop models project global production increases driven primarily by yield increases at high latitudes, even without changes in cropping patterns to match shifts in areas likely to be most productive.

Projected changes in yield for full irrigation are qualitatively and quantitatively similar to those for no irrigation across latitudes (Fig. S6). PEPIC shows substantially greater yield increases in the southern hemisphere for several crops. In certain cases, yields decline more at lower latitudes with full irrigation than with no irrigation. This is because irrigation enables cultivation in otherwise-unsuitable land for these crops, in these models. In summary, both irrigated and unirrigated crop model projections suggest positive latitudinal shifts crop yields over the next half century ^14,15^.

### Projected infection risk changes

Could these yield increases be offset by changing crop disease risk? Infection of plants by pathogens occurs at different rates dependent upon temperature, and each pathogen has a different optimum temperature at which infection of the host is most rapid ^2^. Infection rates are commonly estimated by quantifying the appearance of disease lesions on host plants under controlled conditions ^16^. We estimated relative temperature-dependent infection rates, r(*T*), of 80 fungal and oomycete plant pathogens, for which minimum (*T*_min_), optimum (*T*_opt_) and maximum (*T*_max_) infection temperatures were available in the literature ^2^ (Fig. 1, Table S1). These rates are relative (bound between zero and one) to enable comparison among pathogens. The rate is greatest, i.e. r(*T*) = 1, at *T*_opt_, and declines to zero as temperature decreases to *T*_min_ or increases to *T*_max_. We chose to model infection temperature responses rather than the more commonly-measured growth in culture, because *in planta* responses differ substantially from *in vitro* responses ^2^. Essentially, the temperature range for infection is narrower, and optimum temperature lower, than for growth in culture. However, for two important pathogens, *Magnaporthe oryzae* (causing rice blast) and *Zymoseptoria tritici* (Septoria tritici blotch of wheat), infection temperatures were not available therefore we used lesion development and growth in culture temperatures, respectively. Optimum infection temperatures varied from 10.5 to 34.7 °C among species (median 21.9, IQR 19.6 - 25.0). As global temperatures rise (Fig. S7), infection risks (and distributions) of these pathogens should shift latitudinally ^3^.

**Fig 1.**
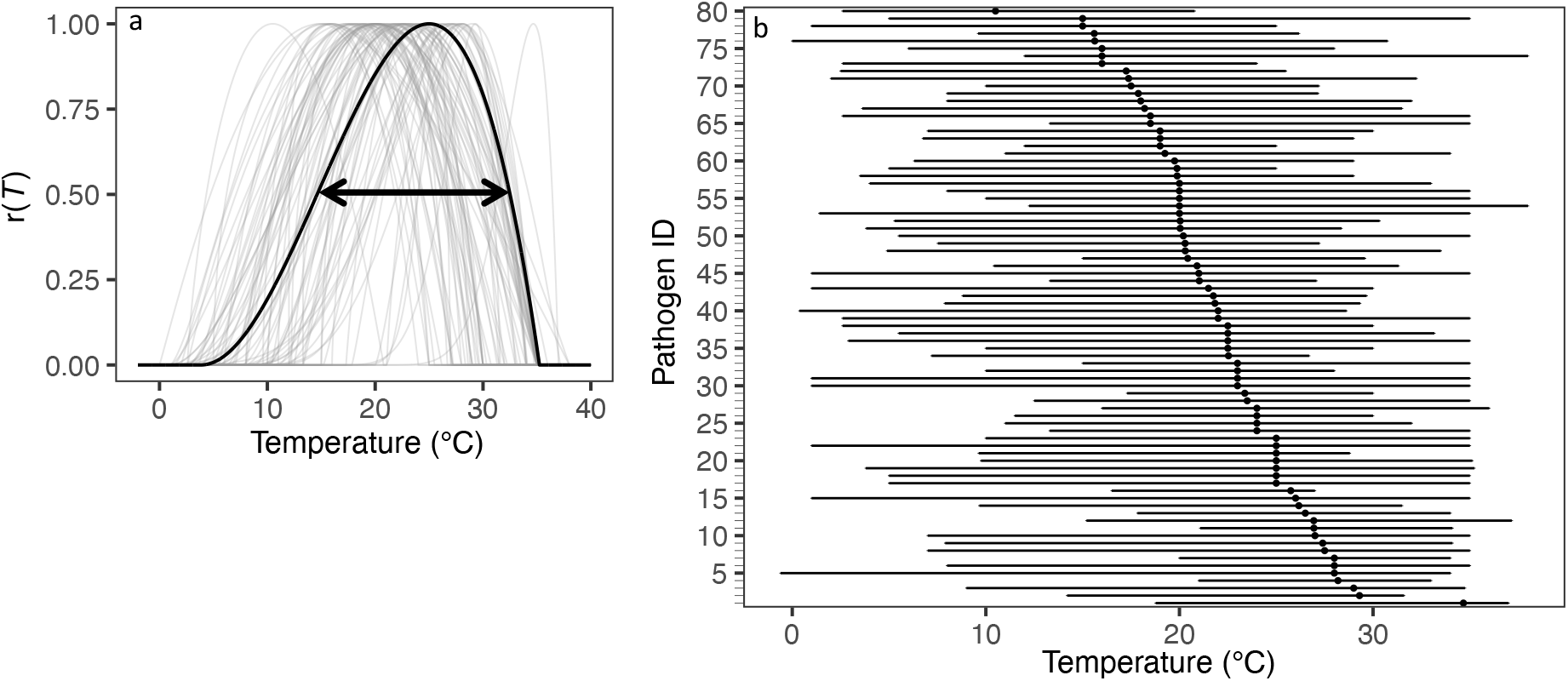
Summary infection cardinal temperature for 80 plant pathogens included in this study. (a) Temperature response curves for r(*T*) determined by *T*_min_, *T*_opt_, and *T*_max_, as well as Equation S1. Arrow indicates temperatures where r(*T*) = 0.5 for an example pathogen. (b) Points refer to *T*_opt_, bars refer to temperature range (defined by *T*_min_ and *T*_max_). Pathogens are ordered by *T*_opt_. Pathogen ID in Table S1.

Defining pathogen species richness, R_r_, as the number of pathogens with r(*T*) ≥ 0.5 for their hosts in a particular location (Figs. S8, S9) at a particular time, we found that R_r_ decreases at low latitudes, and increases at high latitudes, by the end of the 21^st^ Century under RCP 6.0 (Fig. 2a,b). R_r_ increases substantially in Europe and China, but declines in Brazil, sub-Saharan Africa, India and Southeast Asia. Rapid global dissemination by international trade and transport ^17^ means that pathogens are likely to reach all suitable areas that are not yet affected (Fig. S10).

**Fig 2.**
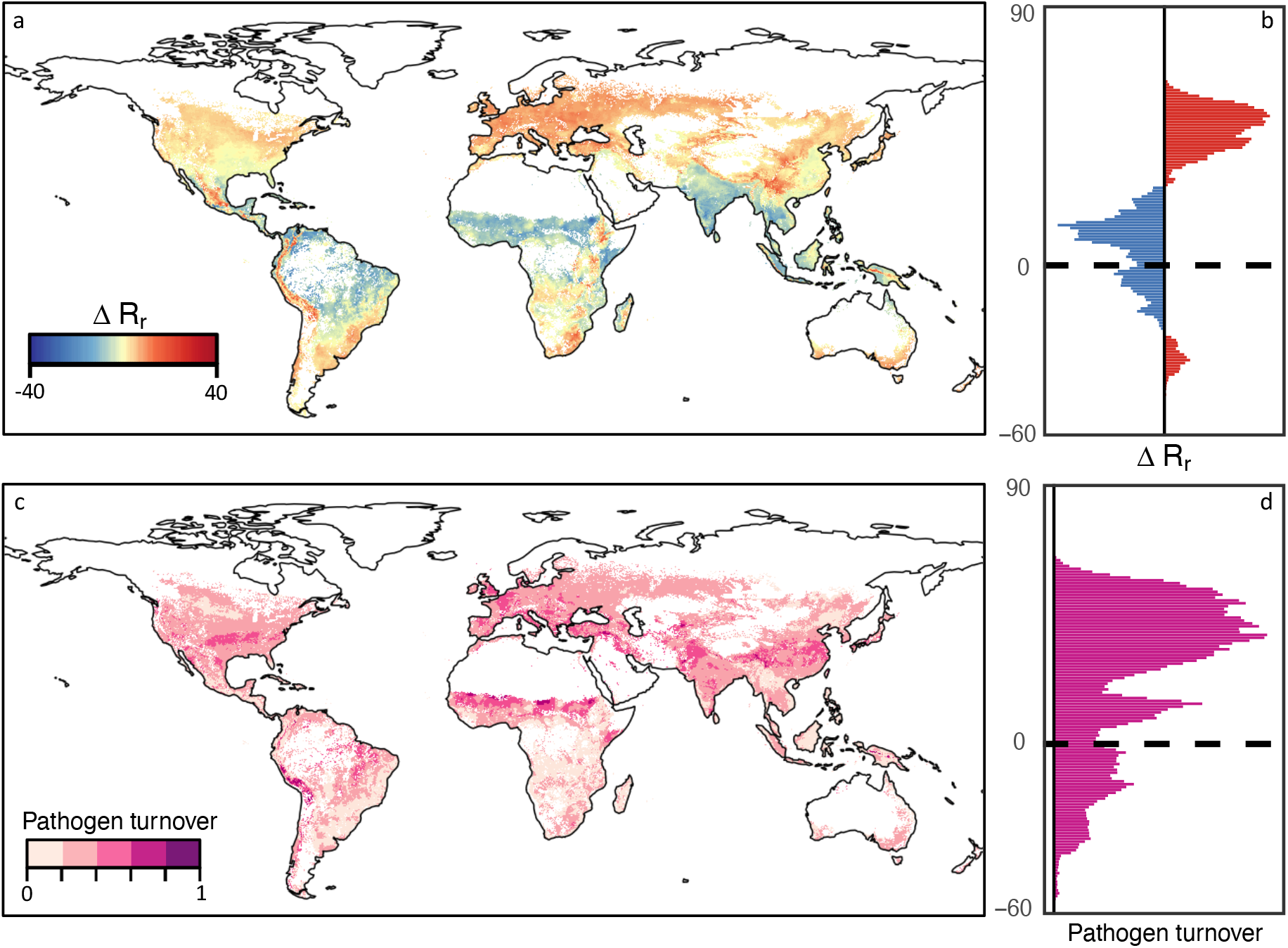
Average change in (a, b) R_r_ and (c, d) pathogen turnover under RCP 6.0 across all months. Red and blue indicate increases and decreases in R_r_, respectively. Darker pink indicates larger changes in pathogen turnover. Pathogens restricted by host distributions extracted from EarthStat. White grid cells contain no hosts and were excluded from the analysis. (b,d) Data aggregated to 1° resolution for plotting.

In our model R_r_ was projected to vary through the year, with the largest increases in North America, Europe and China during northern-Hemisphere autumn (Figs. 3, S11). Decreases in R_r_ are projected at low to mid latitudes in northern-Hemisphere winter, shifting northwards into higher latitudes during summer. India is expected to see large declines in R_r_ over much of the year, with increases in northern parts of India only in winter. Under increasingly strong greenhouse gas emissions scenarios, the overall latitudinal patterns of R_r_ and resultant compositional change in both Hemispheres remain the same, but their amplitudes increase (Fig. 4). R_r_ declined at low latitudes and increased at high latitudes, while compositional changes peaks at around 10° and 30-40°.

**Fig 3.**
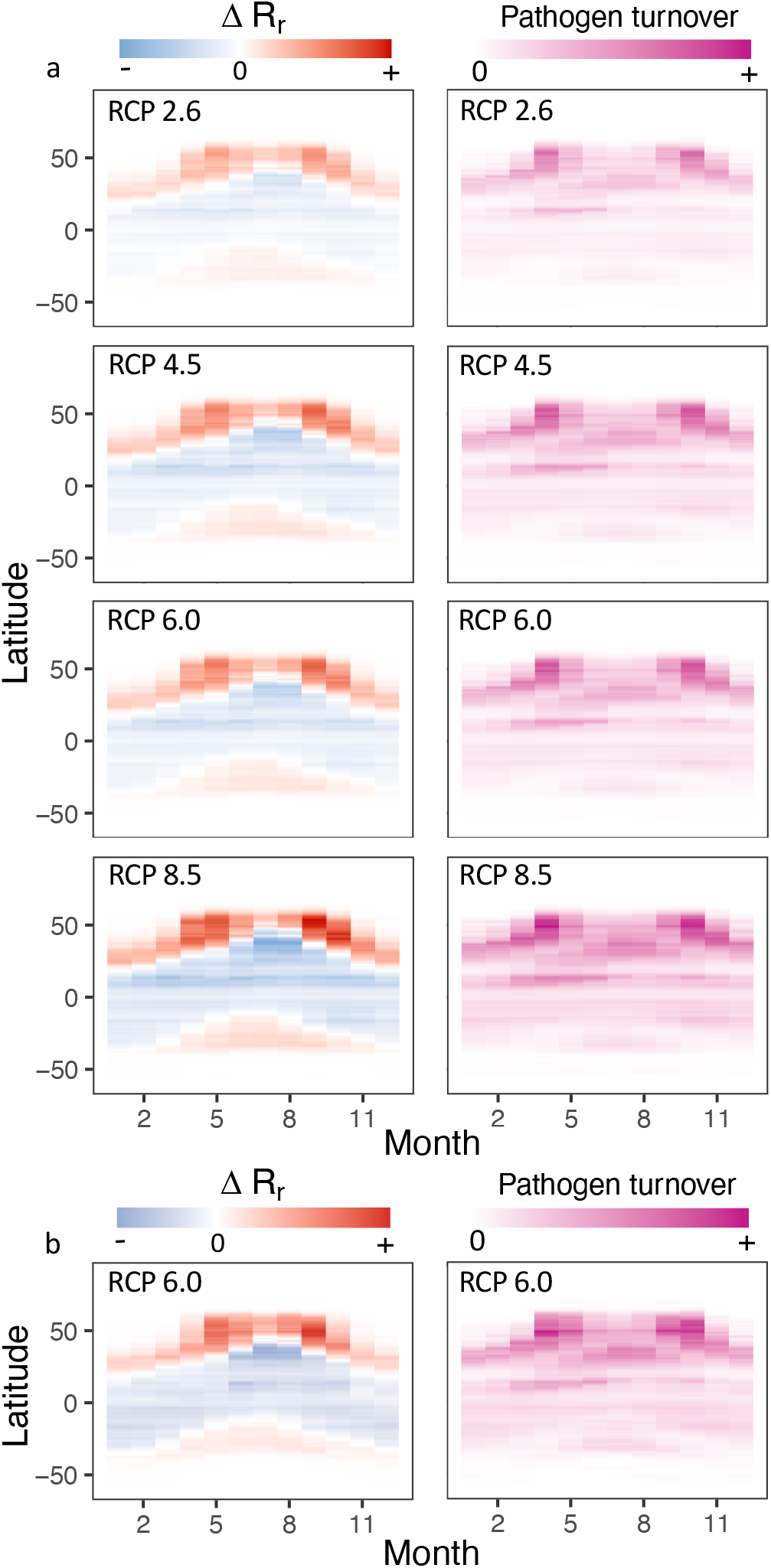
Impact of RCP and pathogen restriction method on change in R_r_ and pathogen turnover. Red and blue indicate increases and decreases in R_r_, respectively. Darker pink indicates larger changes in pathogen turnover. Pathogens restricted by estimates of host distributions extracted from (a) EarthStat and (b) MIRCA2000. Crop calendars only considered in MIRCA2000. Fewer pathogens included in (b) due to fewer host distributions available. Data aggregated to 1° resolution for plotting.

**Fig. 4.**
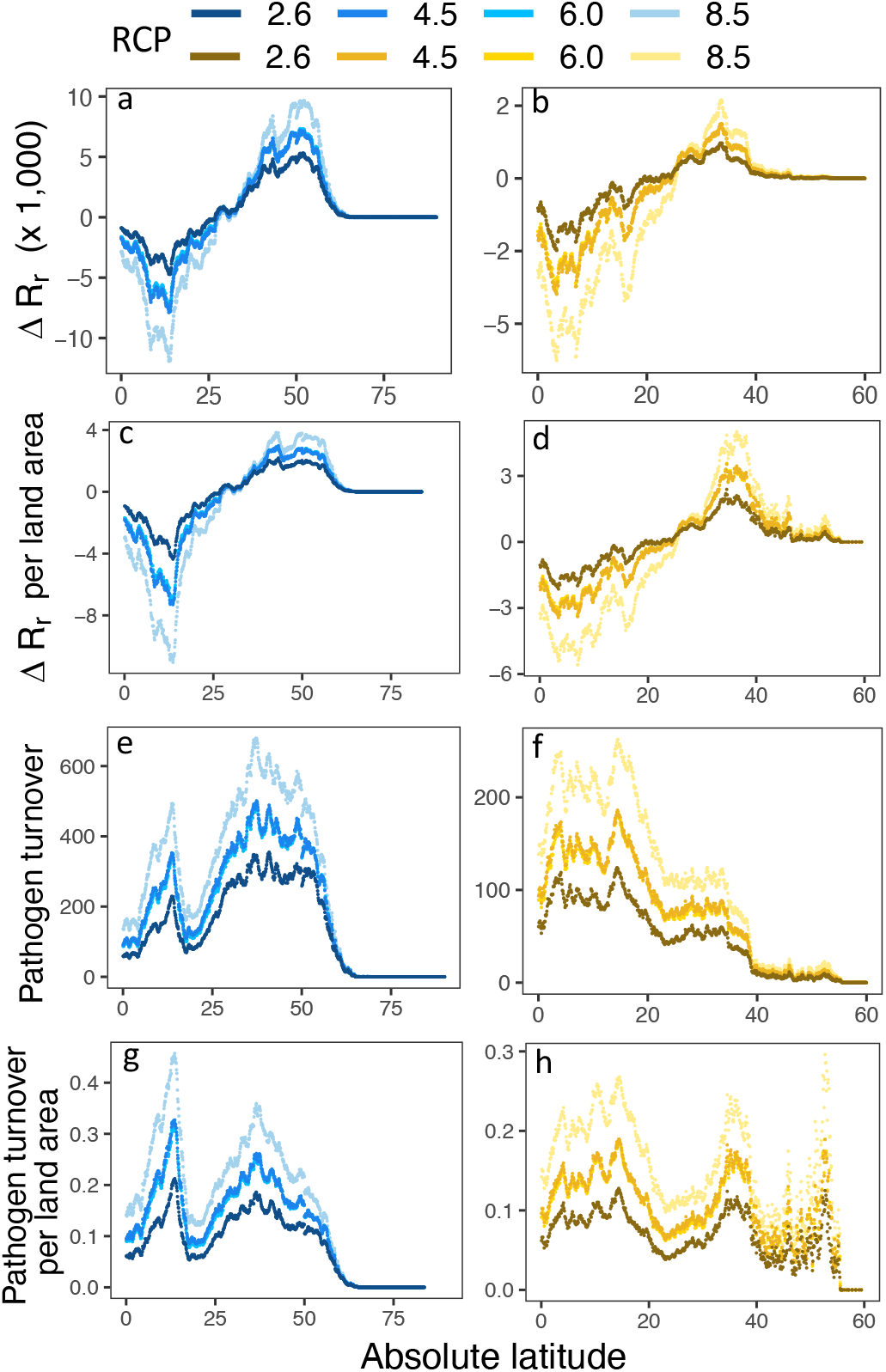
Impact of RCP on average change of R_r_ and pathogen turnover across all months. (a, c, e, g) Northern hemisphere. (b, d, f, h) Southern Hemisphere. Pathogens restricted by estimates of host distributions extracted from EarthStat. Land area refers to total land area, not crop area.

Future changes in pathogen r(*T*) follow changes in yield by latitude for the majority of crops (Fig. 5). The majority of rice pathogens in our sample show increased r(*T*) across all latitudes, with few showing a widespread decline in the tropics. While r(*T*) of several maize pathogens is expected to increase at low latitudes, the risk from many others will decline. Maize, millet and sugarcane are expected to undergo yield declines at low latitudes, but these will be accompanied by declines in r(*T*) from many of their pathogens. Soybean, sunflower and wheat show little yield gain in the tropics, while experiencing reduced r(*T*) from a number of pathogens. Conversely, both yields and r(*T*) increase strongly with latitude. Cassava r(*T*) generally increases near the equator. Overall, high latitudes will see increasing potential crop yields while simultaneously facing a larger r(*T*) by fungal and oomycete pathogens.

**Fig 5.**
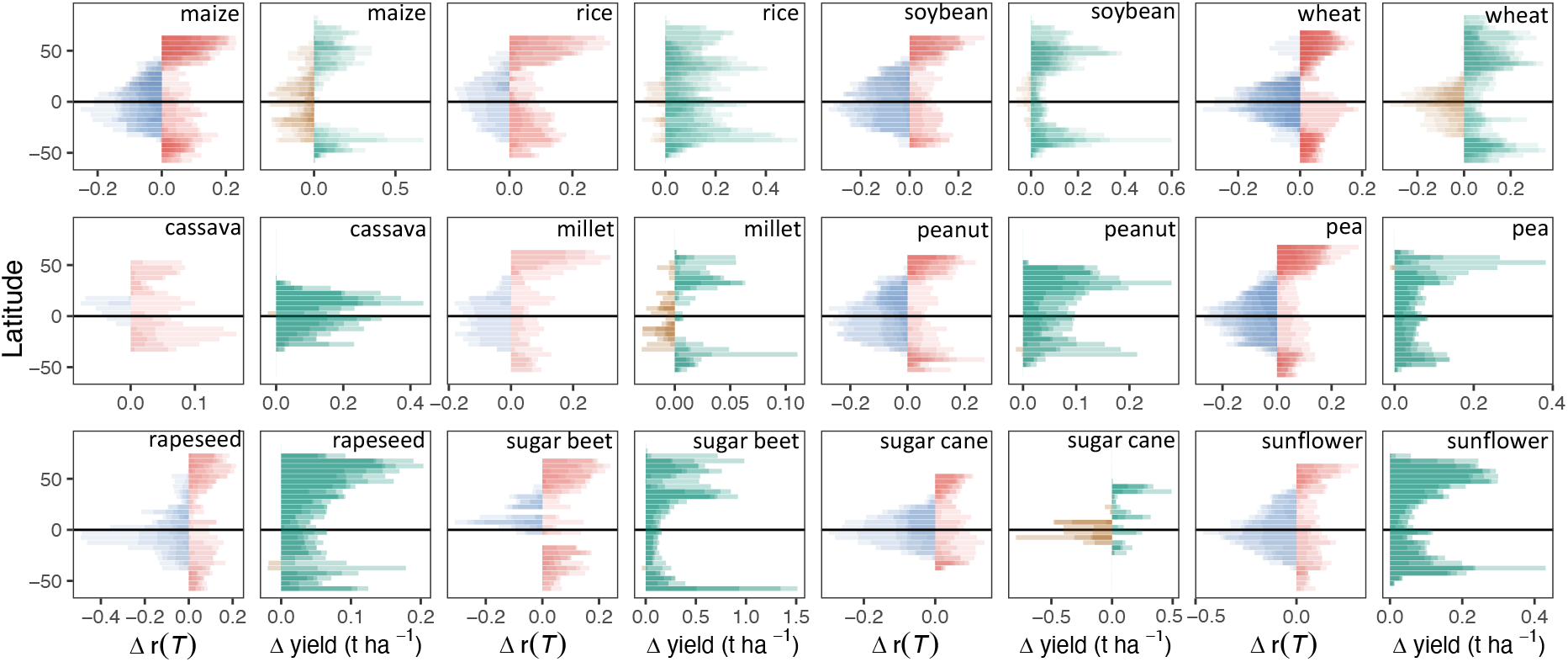
Changes in crop yield and r(*T*) under RCP 6.0 by latitude. Crops are (mai) maize, (ric) rice, (soy) soybean, (whe) wheat, (cas) cassava, (mil) millet, (nut) peanut, (pea) pea, (rap) rapeseed, (sgb) sugar beet, (sug) sugar cane, and (sun) sunflower. Red and blue indicate increases and decreases in r(*T*), respectively. Green and brown indicate increases and decreases in crop yield, respectively, under no irrigation scenario. Colour saturation indicates numbers of pathogens or number of crop- and climate models. Pathogens restricted by estimates of host distributions extracted from MIRCA2000. Data aggregated to 5° for plotting.

We found significant direct spatial matching between future changes in r(*T*) and crop yields (Fig. S12). Correlations between future changes in crop yields and r(*T*) for maize, soybean, sunflower, and wheat exceeded 0.4. Although a weak negative correlation was calculated for cassava (*r* = −0.09), our analysis included far fewer pathogens for this crop, compared to other crops (Table S2). Future crop production, particularly for three major crops, will likely not only be affected directly by climate change, but also indirectly via shifts in plant pathogen distributions.

Changing climate will affect not only the number of pathogens able to infect crops, but also the compositions of pathogen assemblages (Figs. 2cd). Overall, the largest changes in pathogen species composition will occur at high latitudes in the northern Hemisphere, particularly in Europe, China and central to eastern USA. Large changes are also expected in the Sahel, but this region, like much of Brazil, India and southeast Asia, will see declines in overall R_r_. Hence, the change in pathogen assemblage in these areas is unlikely to pose a major threat to production. Europe, China and Peru are highlighted as regions where both overall burden and species turnover are greatest. These regions will therefore experience the greatest number of emerging, i.e. novel, pathogen pressure. Through the year, two pulses of pathogen assemblage change are seen at high latitude in the northern Hemisphere, first around April, second around September (Fig. 3). The largest changes in species composition are expected in Spring and Autumn in northern USA and Canada, Europe, and northern China (Figs. 3, S13). The largest changes in the Sahel are seen during April and May, while the largest changes in India are seen during May and June.

We compared our model predictions against current known pathogen distributions (Fig. S10, Table S3). Restricting predicted distributions by host distributions (EarthStat) improved overall model fit, reducing false positive rates (predicting pathogen presence in regions where the pathogen is currently not reported) and increasing true negative rates (predicting pathogen absence in regions where the pathogen is currently unreported). Like other species distributions models, we predicted areas of suitability and therefore potential distributions of species, and did not attempt to reconstruct observed distributions. Pathogens are spreading globally ^17^, observational records suffer from under-reporting ^4^ and dispersal limitation prevents species from occupying all possible suitable environments ^18^. These factors all likely contributed to the high false positive rates (0.47, IQR 0.37 – 0.57) of our model. However, high false positives rates were more likely in countries with low per capita GDP (Fig. S10h), indicating an under-reporting bias in developing countries ^4^. Importantly, our model did not erroneously confine potential pathogen distributions, as false negative rates (predicting pathogen absence in regions where the pathogen is currently reported) were very low (0.01, IQR: 0.01 – 0.03).

## Discussion

Our analyses are limited by the availability of infection temperature responses in the published literature. These are not a random sample of all known fungal and oomycete plant pathogens. Given that the historical research focus on plant pathogens has been in developed countries at high latitudes ^19^, our sample is biased towards pathogens which have evolved to infect hosts optimally in cooler climates (Fig. S14). However, our sample does include pathogens able to infect both tropical and temperate crops (Figs. S8, S9), hence this bias does not preclude conclusions being drawn for tropical pathogens.

Infection of a susceptible primary host is central to disease development, but other processes such as spore dispersal, overwintering and infection of any alternate hosts are also important in pathogen epidemiology. We have modelled infection only, in common with previous studies on climate change effects on plant pathogens ^11,20^, under the assumption that inoculum will be present, either through long-distance dispersal or overwintering^21^.

We did not attempt to model intra-specific variation in temperature response functions, though such variation does exist ^22,23^. However, analysis of historical pathogen distributions indicates that range shifts have occurred in line with expectation suggesting that temperature adaptation is slow in comparison with climate change ^6^. We employed infection temperatures rather than the more commonly-measured growth in axenic culture ^2^, for all but two pathogens which were included because of their importance in agriculture ^1^. The distinction is important because growth in culture has a wider temperature range for most pathogens ^2^, and models based on growth in culture would suggest a wider geographical range than models based upon infection dynamics.

We only considered temperature as a determinant of infection rates. However, infection by many fungal and oomycete plant pathogens is promoted by wet conditions ^24^. Multi-model mean projections to the end of the 21^st^ Century suggest that precipitation will increase significantly in boreal regions and decrease significantly around the Mediterranean, with smaller and less certain changes elsewhere even under a high-emissions scenario ^25^. Thus, there appears to be no major change in hydrology that would alter our overall conclusions on latitudinal shifts in pathogen burden. In addition, historical shifts in species populations have largely been driven by global warming ^6^. Relative humidity (RH) declines may offset the impact of increased pathogen temperature suitability at higher latitudes, particularly across Europe (Fig. S15). Increased plant infection across Europe has been predicted under future climate, where pathogen temperature tolerances and infection wetting period were considered ^11^. RH was not considered in our model due to paucity of data concerning pathogen RH relations, as well as large uncertainties over future global RH projections ^26^. To investigate the consequences of omitting humidity effects on infection risk, we compared results of models utilizing 3-hourly temperature and leaf wetness estimates with those utilizing only 3-hourly temperature and only monthly temperature during the growing season, for two rust pathogens (see Appendix in Supplementary Information). We found that that the monthly temperature models replicated the overall spatiotemporal patterns seen in the 3-hourly temperature and leaf wetness models, and that infection rate estimates were highly correlated among models. Finally, global observations^27^ and field-scale experiments^28^ suggest that temperature is the most important determinant of fungal distributions and activity.

We did not include potential future changes in crop phenology. Warming is expected to extend the growing season of temperate crops by a few days by the end of the 21^st^ Century, while increasing temperatures may reduce the length of the growing season in tropical crops ^29^. As our seasonal modelling was conducted using monthly crop calendars, the influence of altered growing seasons on our results is likely to be small. We did not include potential future changes in crop distributions. The socioeconomic factors leading to changes in future crop distributions are challenging to predict ^30^, and differing future land use scenarios are beyond the scope of the present analysis. The crop yield projections we employed are subject to uncertainty, both due to the parameterization of the crop models themselves and to the future climate change scenarios ^31,32^. However, the global pattern of greater yield increases at higher latitudes is conserved across models, and accords with the latitudinal trends in temperature.

Future crop yields have been modelled using only plant physiological responses to abiotic conditions. We analysed pathogen temperature physiology to understand how indirect, biotic responses to climate change could impact production. We have shown that crop disease burdens could track crop responses, increasing at higher latitudes where climate change is projected to boost yields.

Furthermore, the suite of crop diseases that farmers face in some of the world’s most productive regions will change dramatically. Crop yield losses to pathogens depend on many factors beyond infection, like host resistance and crop protection ^1^. Agriculture must therefore prepare accordingly if any potential benefits of climate change on crop yields are to be realized.

## Author contributions

DB and TC developed the concept, collated the data, conducted the analyses and prepared the figures. DB wrote the manuscript with contributions from TC and SG.

## Data availability statement

Fungal and oomycete cardinal temperature data are available in Dryad (https://doi.org/10.5061/dryad.tqjq2bvw6) and from Magarey, R. D., Sutton, T. B., & Thayer, C. L. (2005). A Simple Generic Infection Model for Foliar Fungal Plant Pathogens. Phytopatholog 95(1), 92–100 https://doi.org/10.1094/PHYTO-95-0092. The annual crop yield projections data used in this study the Inter-Sectoral Model Intercomparison Project (ISIMIP, https://www.isimip.org). Fungal and oomycete host plant data and geographical distributions (the Plantwise database) were used under license for the current study, and are available with permission from CABI, Wallingford, UK. The FAOSTAT commodity list is available from http://www.fao.org. Global gridded climate data and climate projections are available from WorldClim (https://www.worldclim.org). Global gridded crop distribution data used in this study are available from EarthStat (https://www.earthstat.org) and MIRCA2000 (https://www.uni-frankfurt.de/45218031/data_download). Fungal and oomycete names and name disambiguation data were obtained from Species Fungorum (http://www.speciesfungorum.org/) and MycoBank (http://www.mycobank.org/). Annual per capita GDP at purchasing power parity (PPP) data were obtained from the World Bank (https://data.worldbank.org/). CMIP5 single level monthly near surface RH data were obtained from the Climate Data Store (https://cds.climate.copernicus.eu).

Administrative boundaries for maps were obtained from GADM (https://www.gadm.org). Coastal outlines were obtained from package *rworldmap* version 1.3-6 for R version 4.0.1.

## Code availability statement

All analyses were conducted using existing functions for R version 4.0.1. No significant custom code was used. R code used for data manipulation is available from the corresponding author on reasonable request.

## Methods

### Model summary

A workflow detailing data preparation, model construction, model validation against known pathogen distributions, and RH considerations is presented in Fig. S16.

### Crop yields

Annual crop yield projections from 2006-2099 were obtained from the Inter-Sectoral Model Intercomparison Project (ISIMIP, www.isimip.org) in January 2020. The crop models were LPJmL ^10^, GEPIC ^33^ and PEPIC ^34^. LPJmL simulates changes carbon and water cycles due to land use, phenology, seasonal CO2 fluxes and crop production. GEPIC and PEPIC are derived from the EPIC agricultural yield and water quality model ^35^. In EPIC, potential crop yield is simulated from solar radiation, crop parameters, leaf area index and harvest index (the economic yield per unit aboveground biomass). Each of these crop models was driven by four global circulation models: MIROC5 ^36^, HadGEM2-ES ^37^, GFDL-ESM2M ^38^ and IPSL-CM5A-LR ^39^. Annual crop yield estimates under RCP 6.0, with CO2 fertilization effects, and both the ‘no irrigation’ and ‘full irrigation’ scenarios, were obtained for all available crops at 0.5 ° spatial resolution. Fertilizer application rates are modelled at country scale in each model. Irrigation is modelled using estimates of the area equipped for irrigation per grid cell. GEPIC/PEPIC modelled maize, rice, soybean and wheat. LPJmL additionally included cassava, millet, pea, peanut, rapeseed, sugarbeet, sugarcane and sunflower. Yield differences between the 2060 – 2080 mean and 2010 – 2030 mean were calculated per grid cell.

### Climate data

Global estimates of recent (1970 - 2000 average) and future (2061 - 2080 average) average monthly temperature at 5 arc minute spatial resolution were obtained from the WorldClim database (www.worldclim.org) [accessed 5/2019]. For future estimates, all global climate models (GCMs) of Representative Concentration Pathways (RCP) 2.6, 4.5, 6.0 and 8.5 were obtained (Table S4) [accessed 5/2019]. For each RCP-GCM combination, average future monthly temperature was calculated as the mid-point of average maximum and minimum monthly temperature, as no average estimates were available. For each RCP, average monthly temperature was calculated as the mean of all GCMs for that RCP.

### Pathogen dataset construction

Estimates of pathogen infection cardinal temperature were extracted from two sources ^16,40^. Collectively, only pathogens with at least one minimum (*T*_min_), optimum (*T*_opt_), and maximum (*T*_max_) estimate for infection cardinal temperature were included. To aid matching of species between sources, pathogen species names reported in the latter were updated according to the Species Fungorum database (SFD) (www.speciesfungorum.org) [accessed 4/2020] (Table S5). If no information was available on the SFD, Mycobank (www.mycobank.org) was used as an alternative [accessed 4/2020]. Discovery and sanction author(s) of species were not provided in one source ^16^, and are not considered here. Pathogen species names have previously been processed^40^ and so were not altered. Mean *T*_min_, *T*_opt_, and *T*_max_ infection cardinal temperature were calculated for each pathogen (hereafter referred to as the ‘Pathogen dataset’). Pathogens with nonsensical cardinal temperatures (i.e. mean *T*_opt_ > mean *T*_max_) were excluded from the analysis, as it was not possible to calculate temperature response functions for such pathogens. *Magnaporthe oryzae* and *Zymoseptoria tritici* are two of the most destructive pathogens of rice and wheat ^1^, respectively, but infection temperature estimates are unavailable. We therefore included cardinal temperature for lesion development of *M. oryzae* ^41^, and average growth in culture cardinal temperatures for 18 strains of *Z. tritici* ^42^. It was assumed that average cardinal temperature for each pathogen was identical across all hosts, for each respective pathogen.

The Plantwise database (CABI) [accessed 28/10/2013, by permission] was used to estimate host range of each pathogen in the Pathogen dataset. To improve matching of pathogen species names, some names were updated in the Plantwise database, according to the SFD or Mycobank [accessed 4/2020] (Table S7). We also used included host range information provided by ref.^16^. All plant-pathogen interaction records for hosts recorded in EarthStat (http://www.earthstat.org) and MIRCA2000 ^43^ were extracted from the Plantwise database. To enable matching of host species, scientific names were assigned to plant hosts found in EarthStat and MIRCA2000 (Table S6). The FAOSTAT commodity list (http://www.fao.org) was used to aid this process. Pathogens absent from the extracted plant-pathogen interaction dataset were excluded from the Pathogen dataset. Consequently, 80 pathogens were included in the Pathogen dataset and hence included in this study (Fig. 1, Table S1).

### Estimating global distributions of pathogen hosts

Two approaches were used to estimate global host distributions for each pathogen included in the Pathogen dataset. First, for 150 crops (including forage crops, Table S6), global estimates of average fractional proportion grid cell harvested (5 arc minute spatial resolution) were obtained from EarthStat ^44^ (http://www.earthstat.org). Crops that could not be clearly identified as species (e.g. “mixed grain”) or contained a large number of different plant genera (e.g. “vegetables”) were excluded. Most crops classified as “not elsewhere specified” (nes) were also excluded. For 150 crops, each crop map was converted to binary presence/absence. If grid cell harvest area fraction was ≥ 0.00001 (equivalent to 0.1 m^2^ ha^−1^), the host was estimated as present in that grid cell. If < 0.00001, hosts were assumed absent. These values were chosen to ensure that crops were estimated as present in grid cells, even if average fractional proportion harvested were estimated as very small. This approach enabled estimation of global distribution for each crop in EarthStat. The Earthstat crop distribution dataset does not provide crop calendars (i.e. the months during which the crop is growing).

Second, for 22 crops (Table S6), global estimates of growing season periods (around the year 2000) were extracted from MIRCA2000 at 30 arc minute spatial resolution ^43^, and resampled to 5 arc minute resolution using neighbour joining algorithm in package *raster* for R ^45^. For each crop, rainfed and irrigated growing season estimates were combined. This provided global monthly estimates of global host presence (within growing season) and absence (outside of growing season), and hence monthly global distribution estimates, at 5 arc minute spatial resolution for 22 crops.

For each pathogen, global distributions for all recorded hosts were combined, and converted to binary presence/absence. This provided a single potential geographical distribution of each pathogen, based on reported pathogen host range (Plantwise) and geographic host distributions (EarthStat or MIRCA2000) (Fig. S8, S9). For example, if a pathogen was recorded in the Plantwise database to successfully infect four hosts recorded in EarthStat, any grid cells that were estimated to contain ≥ 1 of these hosts were converted to 1 (present), and grid cells that there were estimated to contain 0 hosts were converted to 0 (absent). This was done independently for host distributions estimated from EarthStat and MIRCA2000, resulting in two alternative potential geographical distribution of each pathogen. Where MIRCA2000 was utilised, fewer pathogens were included, due to fewer crop species. Further, where host range was estimated from MIRCA2000, the potential geographical range of a pathogen of estimated each month, due to host growing season (Fig. S9). Host ranges were assumed independent for each pathogen, i.e. competition between pathogens for particular hosts was assumed to not occur.

### Modelling pathogen temperature-dependent infection risk

Relative temperature-dependent infection rates, r(*T*), were calculated by a beta function ^46^ (Equation S1) for each pathogen (Fig. 1, Table S1), for all climate data detailed above. We defined pathogen species richness R_r_ as the number of pathogens with r(*T*) ≥ 0.5, i.e. those pathogens with high predicted infection rates. R_r_ acted as a summary metric of pathogen risk per grid cell.

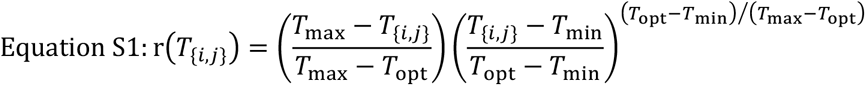

where *i* is the month and *j* is the grid cell.

### Model validation

Pathogen presence (defined as r(*T*) ≥ 0.5) was calculated for recent average monthly temperature estimates utilising two alternative approaches. In the ‘temperature-only model’, pathogens were not restricted by host distributions. In the ‘temperature+host model’, pathogens were additionally restricted by host distributions estimated from EarthStat. In both model iterations, a summary potential global distribution of each pathogen was calculated, whereby if a pathogen was modelled as ‘present’ in a grid cell (*j*) during any month (*i*), then the pathogen was recorded as ‘present’ in that grid cell (*j*).

Outputs from both model iterations were compared to observed records of pathogen presence at country or state scale (hereafter collectively referred to as ‘region’, 396 regions total), from the CABI Plantwise database. Pathogen names in this dataset were updated according to the SFD or Mycobank [accessed 4/2020] to improve matching to the Pathogen dataset (Table S7). Discovery and sanction author(s) of species were not provided in this dataset, and so were not considered here. Thirteen pathogens (*Alternaria cucumerina*, *Botrytis cinerea*, *Cercospora carotae*, *Didymella arachidicola*, *Diplocarpon earlianum*, *Fusarium oxysporum f.sp. conglutinans*, *Fusarium roseum*, *Globisporangium ultimum*, *Nothopassalora personata*, *Puccinia menthae*, *Septoria glycines*, *Stigmina carpophila*, and *Wilsoniana occidentalis*) were excluded from model validation, due to an apparent lack of observational records.

Models were run at 5 arc minute resolution, whereas observed pathogen records were at regional scale (Fig. S10a, c). Hence, model outputs were summed to regional scale (Fig. S10b, d). If a pathogen was modelled as ‘present’ in any grid cell (*j*) in a region, for any month (*i*), the pathogen was modelled as ‘present’ at the regional scale. Gross domestic product based on purchasing power parity (GDP (PPP)) and research output (number of publications) were obtained from the World Bank Data website for 230 territories (data.worldbank.org) [accessed 11/2018]. For the temperature+host model, for each pathogen, median GDP (PPP) and median research output were calculated for territories where (1) both the temperature+host model estimated, and the Plantwise database recorded a pathogen as present (true positive (Sensitivity)), and where (2) the temperature+host model estimated a pathogen as present, but the Plantwise database recorded a pathogen as absent (false positive (Type 1 error)). Data were compared by Welch’s Two Sample two-tailed *t*-test. Where GDP (PPP) and research output were recorded at country scale, but pathogen records were recorded at state scale, states were assigned country-level GDP (PPP) and research output.

### Changes in global temperature-dependent infection risk

We calculated R_r_ for recent and future average monthly (*i*) grid cell (*j*) temperature (*T*_{*i,j*}_), utilising two alternative host-restriction approaches. First, pathogens were restricted by host distributions estimated from EarthStat, for each future climate scenarios (RCP 2.6, 4.5, 6.0, and 8.5). Second, pathogens were restricted by host distributions estimated from MIRCA2000, and RCP 6.0 was used to estimate future average monthly temperature. This allowed for comparison between host restriction method on model outputs of change in spatial patterns of R_r_.

For each model, change in R_r_ was calculated as R_r_ under future climate conditions, minus R_r_ under recent climate, for each grid cell (*j*), for each month (*i*). Within a grid cell, increases or decreases in R_r_ do not reflect the change of species composition ^7^. Therefore, for each model, a modified Jaccard (*J*) index (1 - *J*) of community dissimilarity (pathogen turnover, Equation S2) ^7,47^ was calculated to characterize the change in community composition in each grid cell (*j*), for each month (*i*). High pathogen turnover indicates high community dissimilarity or a large change in species composition.

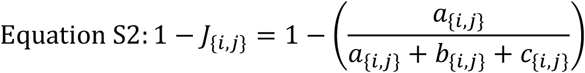

where *a* is the number of pathogens common to a grid cell under recent and future climate, *b* is the number of pathogens unique to a grid cell under recent climate, *c* is the number of pathogens unique to a grid cell under future climate, *i* is the month and *j* is the grid cell. Pathogen turnover was defined as zero for grid cells with no pathogens under both recent and future climates.

### Comparison between future changes in crop yields and r(T) by latitude

For each pathogen of each crop included in MIRCA2000, change in r(*T*) between current and future climate (RCP 6.0) was calculated for each grid cell (*j*), for each month (*i*) (Table S2 provides the number of pathogens included for each crop). Pathogens were restricted by crop distributions estimated from MIRCA2000 (see above). For this analysis, we used estimates from MIRCA2000 for pulses as a proxy for pea crop (*Pisum sativum*). For each crop-pathogen combination, mean change in r(*T*) was calculated for each latitude (5 arc minute resolution), and then aggregated to 5° resolution for plotting using the aggregate function in package *raster* for R ^45^ (Fig. 5).

We tested for evidence of spatial matching between projected changes in crop yield and pathogen r(*T*). For each crop, Pearson correlations (*r*) and spatial cross-correlations (*r*c) were calculated between overall mean change in crop yield and pathogen r(*T*), aggregated to 2° resolution. In this case, compared overall mean change in r(*T*) for all months, for all pathogens with overall mean change in yield from all available models under the no irrigation scenario. Spatial cross-correlations were calculated using the package *spatialEco* for R ^48^. An inverse power law transformation was performed to derive a spatial weights matrix in the analysis of each crop.

### Pathogen sampling bias

Northern and southern latitudinal ranges for plant pests and pathogens were extracted from the CABI Plantwise database. As previous, some pathogen names in this dataset were updated according to the SFD or Mycobank [accessed 4/2020] to improve matching to the Pathogen dataset (Table S7) and 13 pathogens were excluded from the analysis, due to an apparent lack of observational records. Pathogen names were not updated in this dataset if they were absent from the Pathogen dataset. Northern and southern latitudinal ranges for pathogens included in the Pathogen dataset were compared to that of all fungi and oomycetes pathogens for which latitudinal ranges were available.

### Relative humidity considerations

Coupled Model Intercomparison Project 5 (CMIP5) single level monthly near surface RH data (0.125° to 5° spatial resolution depending on model) were extracted from the Climate Data Store (https://cds.climate.copernicus.eu). Data from all available future (RCP 6.0, 2070) and corresponding recent (1985) model-ensemble combinations (see Table S8 for further details) were extracted from NC files and converted to *raster* objects.

For each model-ensemble-month combination, change in RH was calculated as future RH minus recent RH. If a model had multiple ensembles, mean change for each month was calculated from all ensembles. All data were resampled to 5 arc minute resolution using bilinear algorithm in package *raster* for R. Mean monthly change in RH was calculated from all model estimates to provide single monthly estimates. Grid cells that contained no hosts in the EarthStat database were excluded from the analysis. Hence, only grid cells included in analyses of R_r_ and pathogen turnover were included. Grid cells were aggregated to 2° spatial resolution to calculate Pearson correlations (*r*) between change in RH and change in R_r_ (RCP 6.0) for March, June, September, and December.

The Appendix (see Supplementary Information) compares r(*T*) estimates from models using 3-hourly temperatures estimates constrained by leaf wetness, with results obtained using only monthly average temperatures unconstrained by leaf wetness.

