## Supplementary Information for "Plant pathogen infection risk tracks global crop yields under climate change"

Supplementary Figures (Fig. S1 – S16)

Supplementary Tables (Table S1 – S8)

Appendix - The influence of canopy moisture on estimates of temperature-dependent infection risk

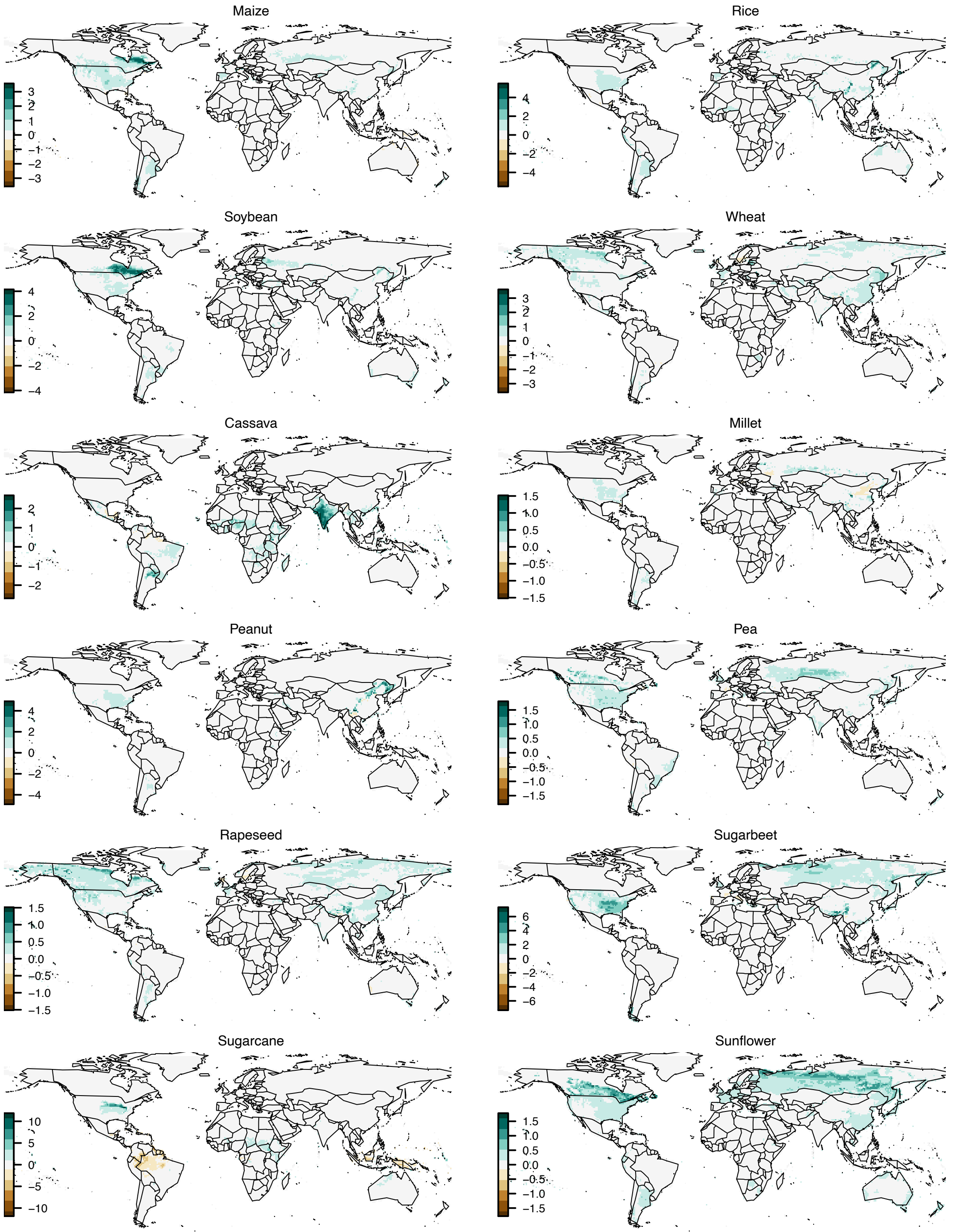

**Fig. S1. Projected yield differences (2020 – 2070), LPJmL crop model**. Values are difference between 2061 – 2080 mean and 2011 – 2030 mean (t $ha^{-1}$), averaged over four climate models (GFDL-ESM2M, HADGEM2-ES, IPSL-CM5A-LR, MIROC5).

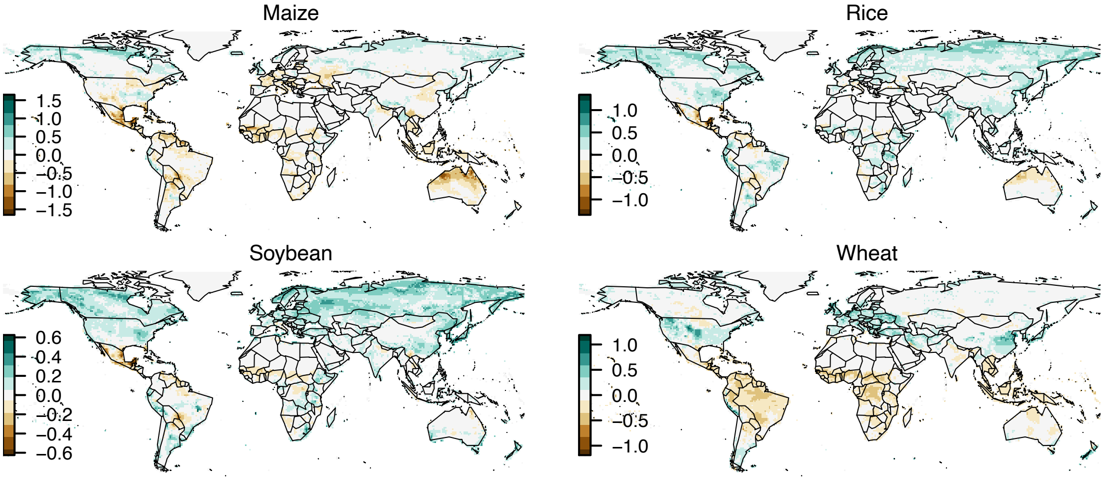

**Fig. S2. Projected yield differences (2020 – 2070), GEPIC crop model**. Values are difference between 2061 – 2080 mean and 2011 – 2030 mean (t ha$^{-1}$), averaged over four climate models (GFDL-ESM2M, HADGEM2-ES, IPSL-CM5A-LR, MIROC5).

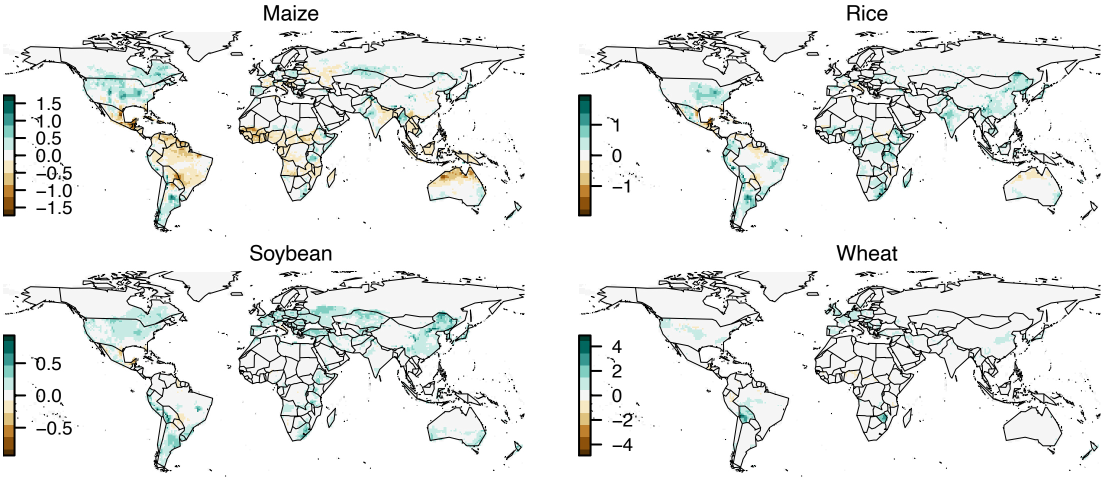

**Fig. S3. Projected yield differences (2020 – 2070), PEPIC crop model**.

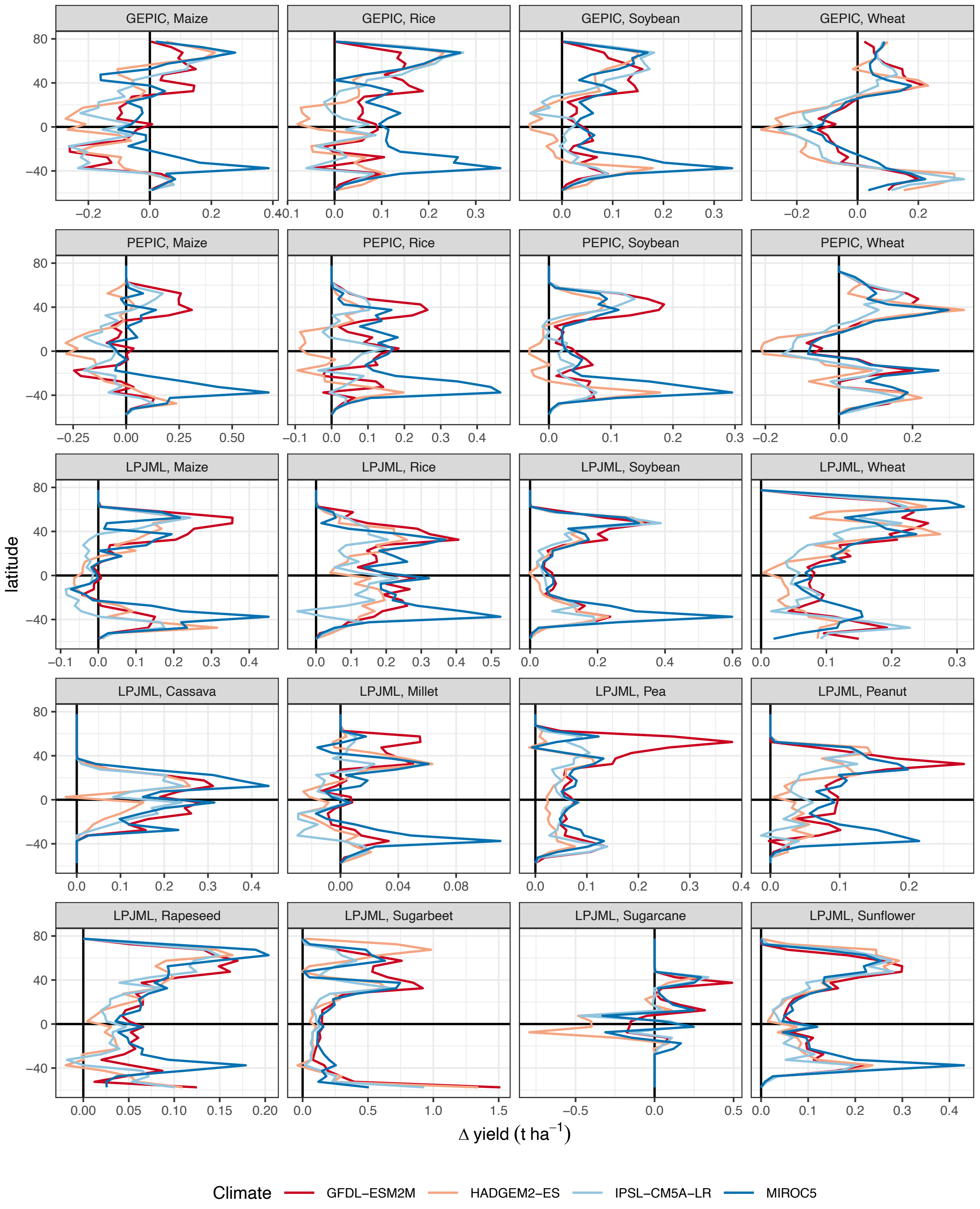

**Fig. S4. Projected yield differences (2020 – 2070) by latitude, per crop model and crop**. Data are averaged over 5° latitudes. Major crops are in the first three rows.

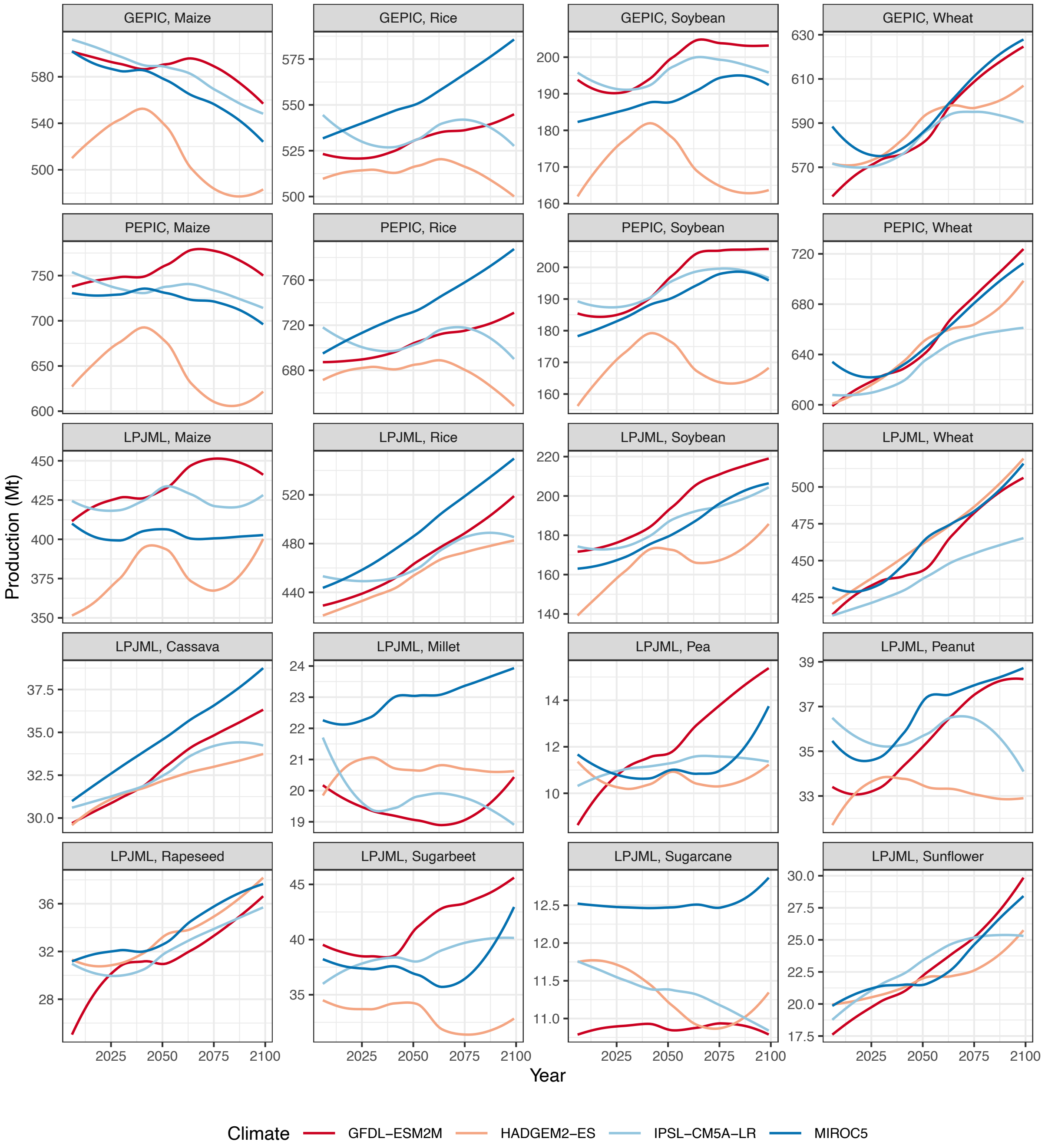

**Fig. S5. Projected global production (2006 – 2099) by crop model and crop, for four climate models**. Production estimates derived by product of projected yields and crop harvested area in year 2000 at 0.5° resolution. Lines show local polynomial regression (loess) smooths of the data. Changes in crop distributions were not considered.

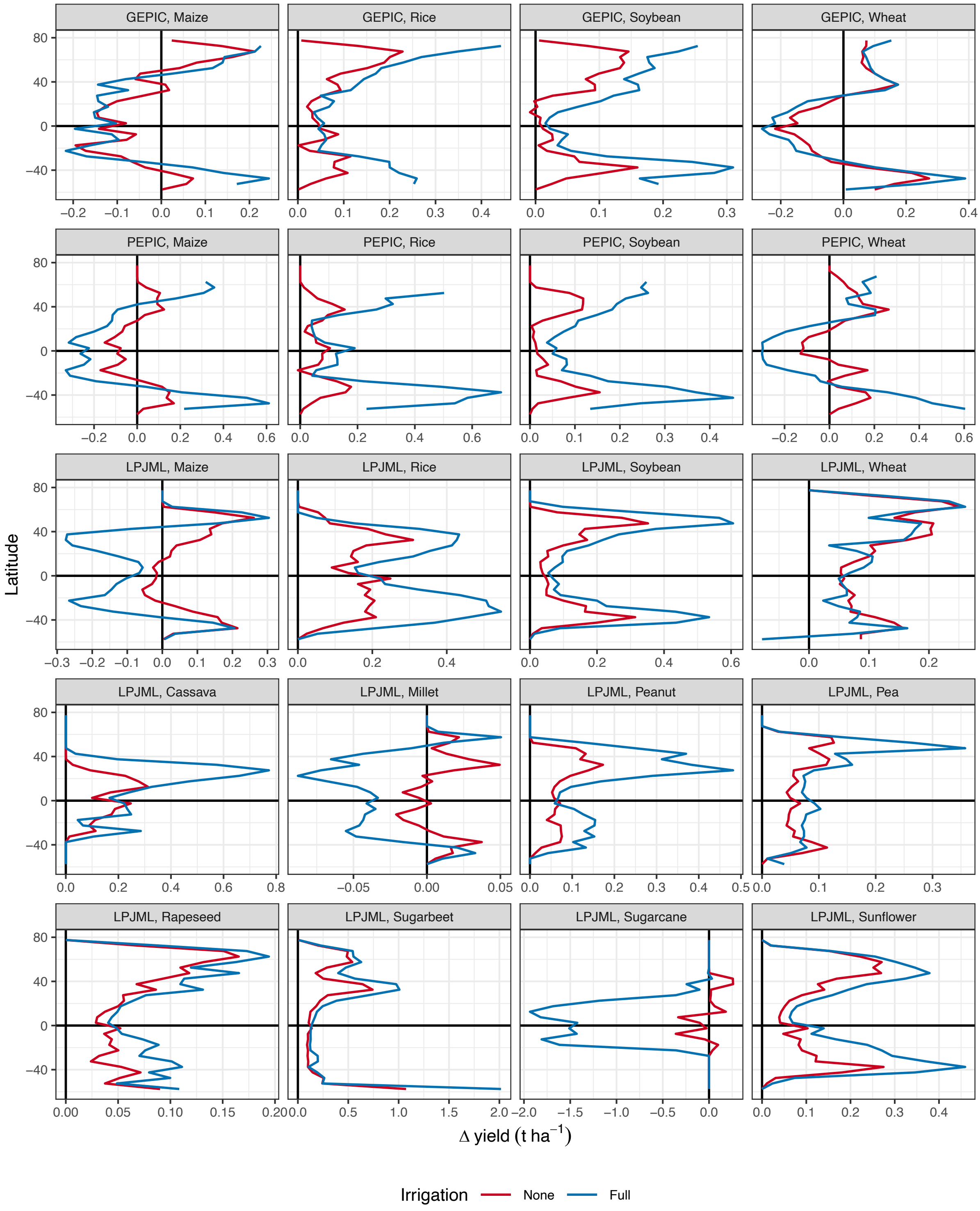

**Fig. S6. Projected yield differences (2020 – 2070) by latitude, per crop model, crop and irrigation scenario**. Data are averaged over 5° latitudes. Major crops are in the first three rows. Lines show means of projections for each of the four climate models.

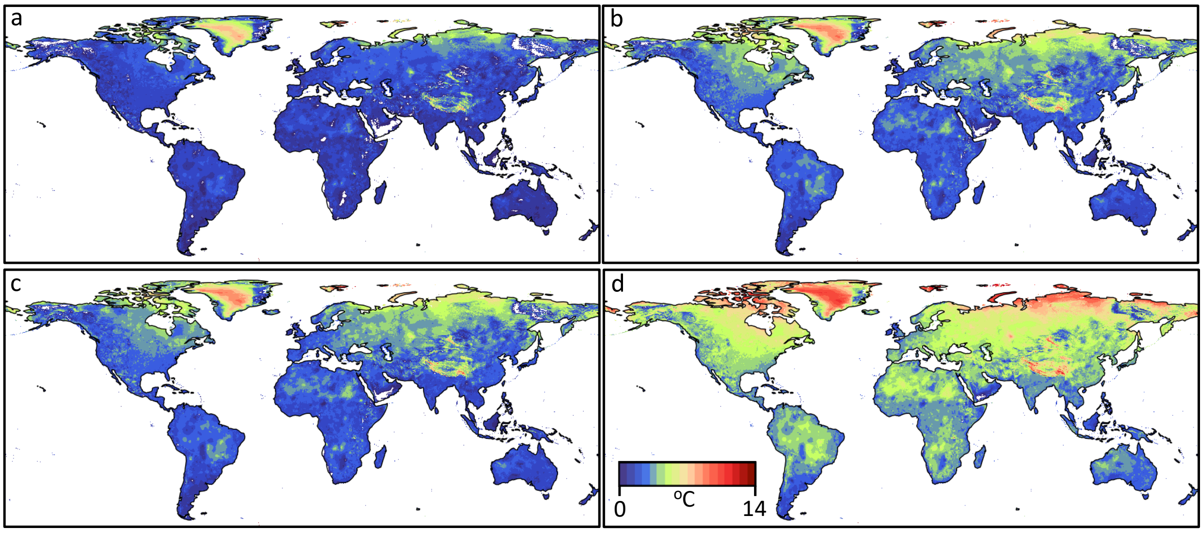

**Fig S7. Average increase in temperature between recent (1970 - 2000 average) and predicted future (2061 - 2080 average) climate under four Representative Concentration Pathways (RCP) scenarios**. RCP (a) 2.6, (b) 4.5, (c) 6.0, (d) 8.5. For clarity, temperature changes ≤ 0 ºC and > 14 °C are excluded.

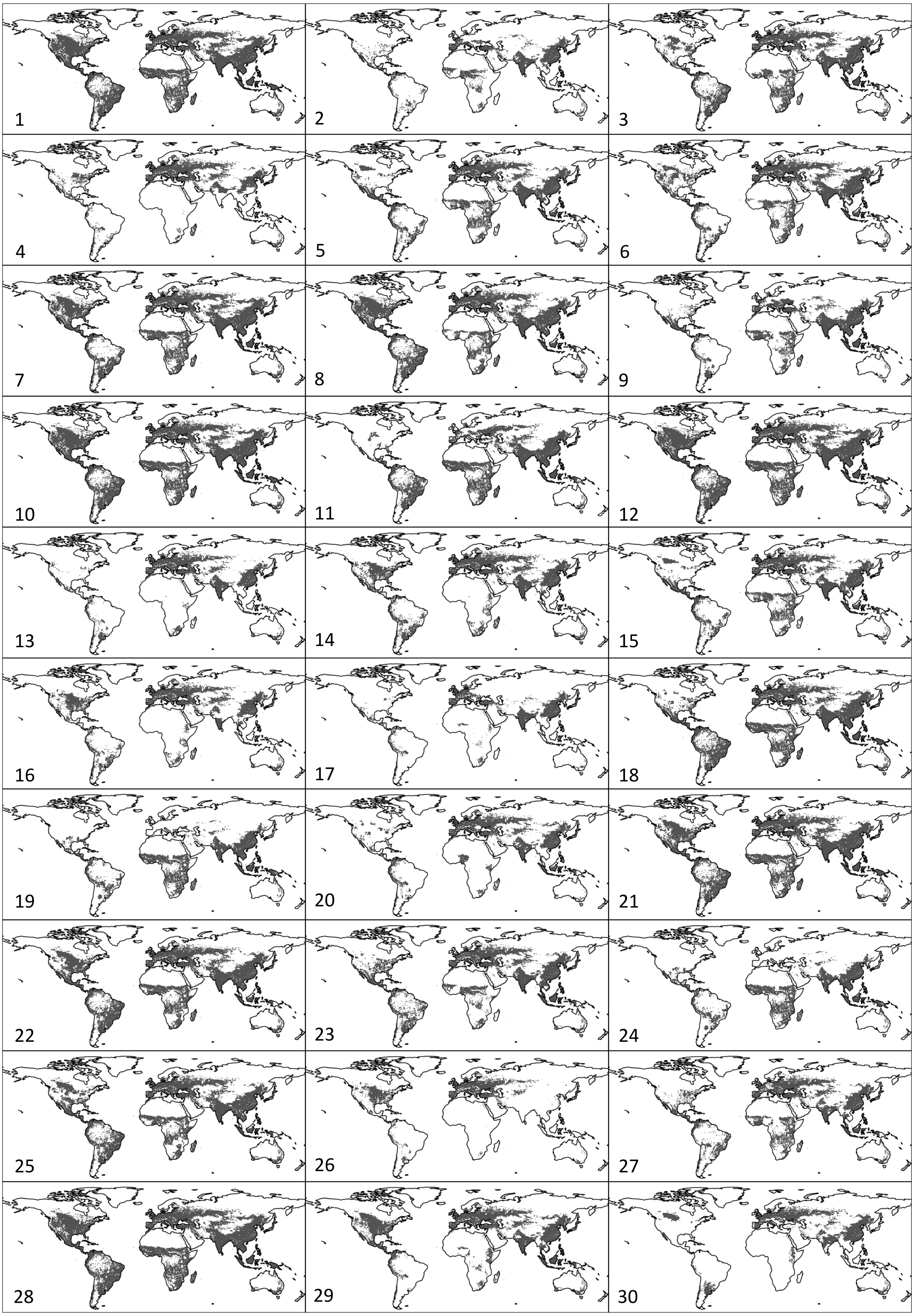

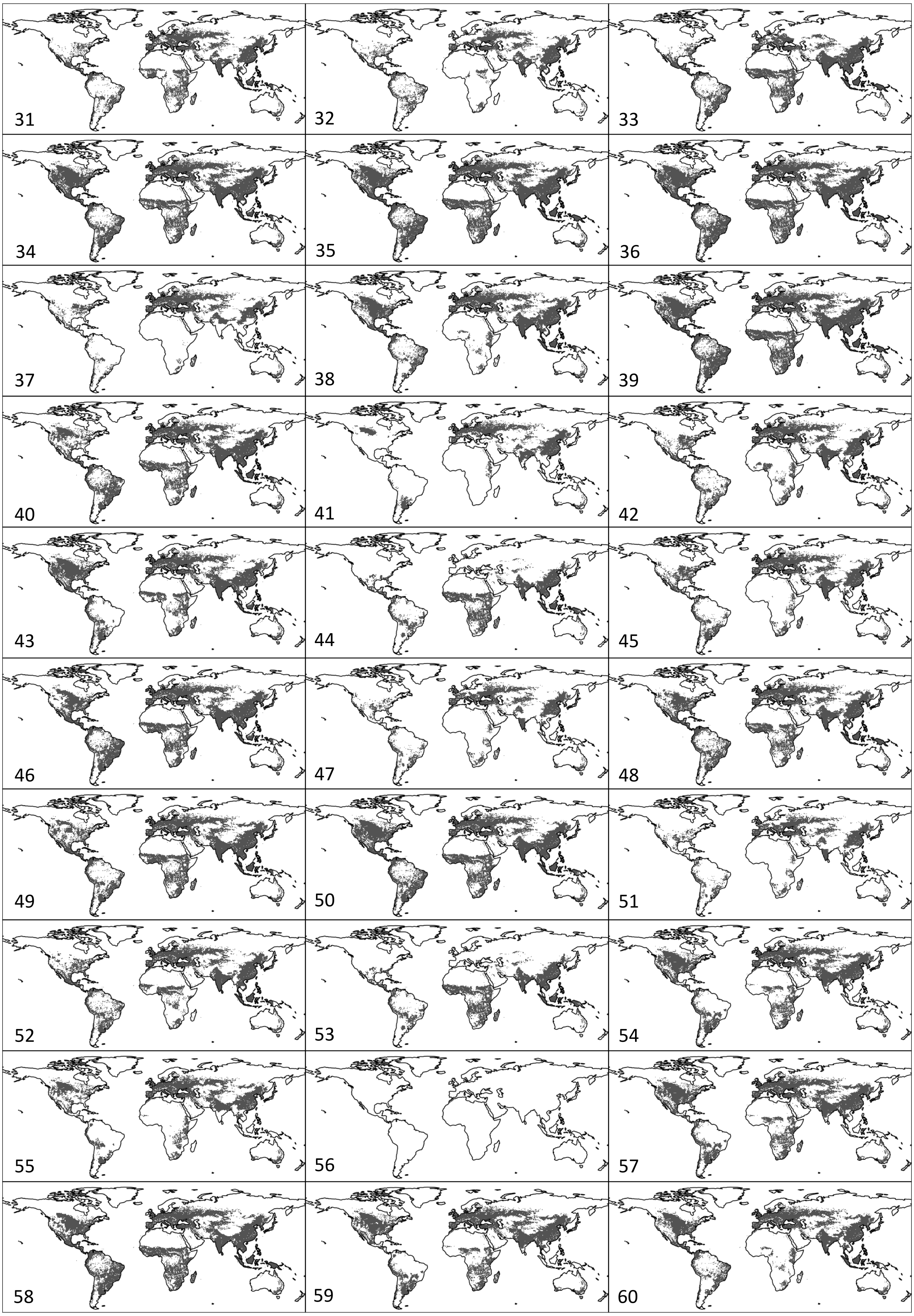

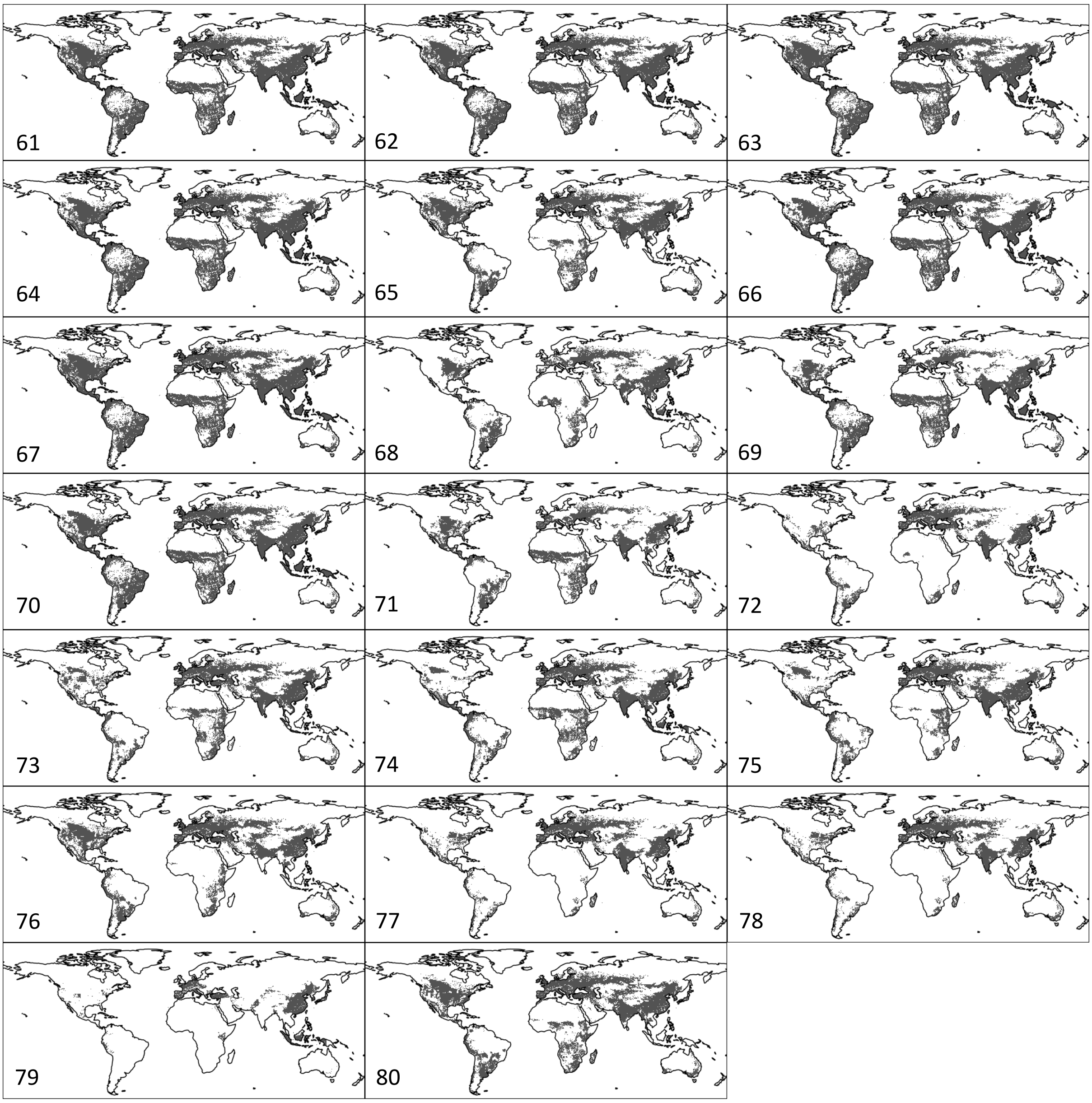

**Fig S8. Binary host distributions estimated from EarthStat**. In cases where a pathogen was recorded in the Plantwise database as being able to infect > 1 hosts recorded in EarthStat, host distributions were combined and converted to binary presence/absence. Grey refers to grid cells that contain at least 1 host. White refers to grid cells that contain no hosts. (1) *A. brassicae*, (2) *A. cucumerina*, (3) *A. longipes*, (4) *A. mali*, (5) *A. porri*, (6) *A. radicina*, (7) *A. solani*, (8) *A. euteiches*, (9) *A. psidii*, (10) *B. basicola*, (11) *B. oryzae*, (12) *B. sorokiniana*, (13) *B. jaapii*, (14) *B. dothidea*, (15) *B. squamosa*, (16) *B. cinerea*, (17) *B. lactucae*, (18) *C. fimbriata*, (19) *C. arachidicola*, (20) *C. carotae*, (21) *C. acutatum*, (22) *C. lindemuthianum*, (23) *C. orbiculare*, (24) *D. arachidicola*, (25) *D. pinodes*, (26) *D. earlianum*, (27) *F. fulva*, (28) *F. graminearum*, (29) *F. oxysporum f.sp. conglutinans*, (30) *F. oxysporum f.sp. lini*, (31) *F. oxysporum f.sp. lycopersici*, (32) *F. oxysporum f.sp. niveum*, (33) *F. oxysporum f.sp. vasinfectum*, (34) *F. roseum*, (35) *G. debaryanum*, (36) *G. ultimum*, (37) *G. juniperi-virginianae*, (38) *L. maculans*, (39) *M. phaseolina*, (40) *M. oryzae*, (41) *M. lini*, (42) *M. fructicola*, (43) *M. rabiei*, (44) *N. personata*, (45) *P. obtusa*, (46) *P. pachyrhizi*, (47) *P. ampelicida*, (48) *P. cactorum*, (49) *P. infestans*, (50) *P. nicotianae*, (51) *P. viticola*, (52) *P. cubensis*, (53) *P. arachidis*, (54) *P. graminis*, (55) *P. hordei*, (56) *P. menthae*, (57) *P. recondita*, (58) *P. sorghi*, (59) *P. striiformis*, (60) *P. brassicae*, (61) *P. teres*, (62) *P. arrhenomanes*, (63) *R. solani*, (64) *R. stolonifer*, (65) *R. secalis*, (66) *S. graminicola*, (67) *S. sclerotiorum*, (68) *S. glycines*, (69) *S. cruentum*, (70) *S. reilianum*, (71) *S. sorghi*, (72) *S. carpophila*, (73) *S. endobioticum*, (74) *U. cepulae*, (75) *U. viciae-fabae*, (76) *U. avenae*, (77) *V. inaequalis*, (78) *V. pyrina*, (79) *W. occidentalis*, (80) *Z. tritici*.

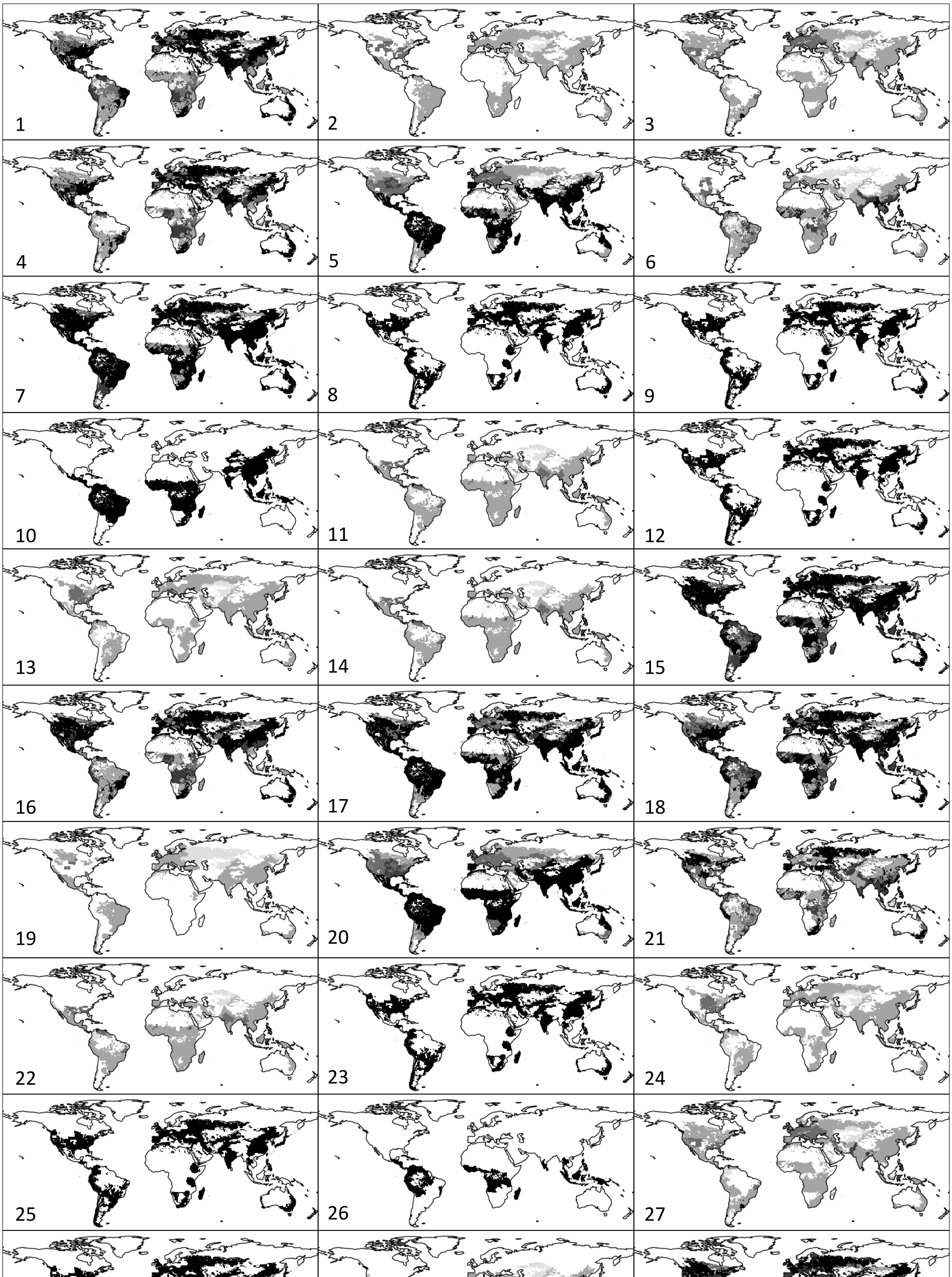

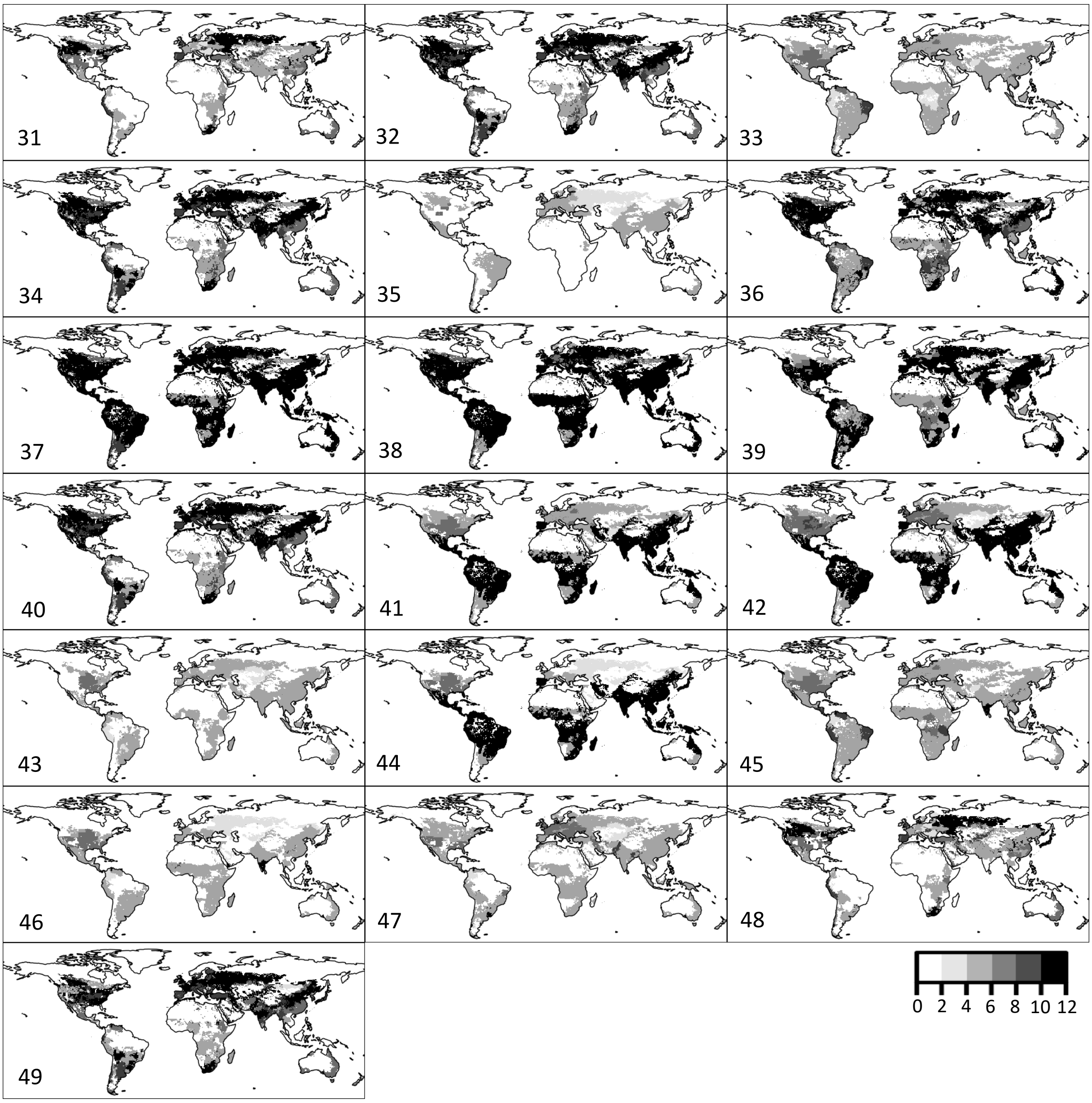

**Fig S9. Binary host distributions estimated from MIRCA2000**. In cases where a pathogen was recorded in the Plantwise database as being able to infect > 1 hosts recorded in MIRCA2000, host distributions were combined and converted to binary presence/absence. Grey scale refers to the number of months a grid cell was recorded to contain at least one host. (1) *A. brassicae*, (2) *A. longipes*, (3) *A. radicina*, (4) *A. solani*, (5) *B. basicola*, (6) *B. oryzae*, (7) *B. sorokiniana*, (8) *B. dothidea*, (9) *B. cinerea*, (10) *C. fimbriata*, (11) *C. arachidicola*, (12) *C. acutatum*, (13) *C. lindemuthianum*, (14) *D. arachidicola*, (15) *F. graminearum*, (16) *F. roseum*, (17) *G. debaryanum*, (18) *G. ultimum*, (19) *L. maculans*, (20) *M. phaseolina*, (21) *M. oryzae*, (22) *N. personata*, (23) *P. obtusa*, (24) *P. pachyrhizi*, (25) *P. ampelicida*, (26) *P. cactorum*, (27) *P. infestans*, (28) *P. viticola*, (29) *P. arachidis*, (30) *P. graminis*, (31) *P. hordei*, (32) *P. recondita*, (33) *P. sorghi*, (34) *P. striiformis*, (35) *P. brassicae*, (36) *P. teres*, (37) *P. arrhenomanes*, (38) *R. solani*, (39) *R. stolonifer*, (40) *R. secalis*, (41) *S. graminicola*, (42) *S. sclerotiorum*, (43) *S. glycines*, (44) *S. cruentum*, (45) *S. reilianum*, (46) *S. sorghi*, (47) *S. endobioticum*, (48) *U. avenae*, (49) *Z. tritici*.

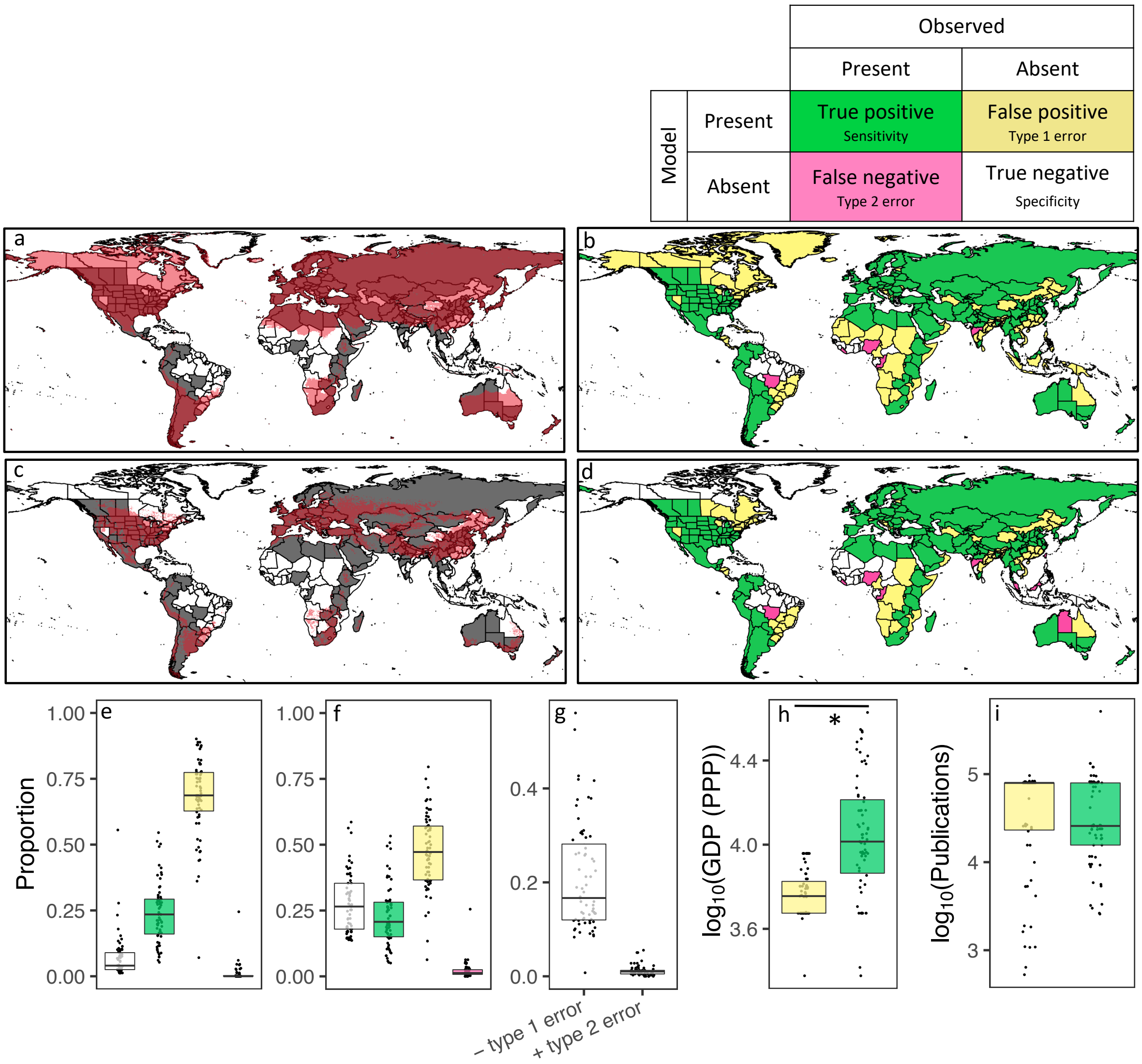

**Fig S10. Model comparison under recent climate conditions**. (a - d) Example model output maps (*P. striiformis*). (a, c) Spatial resolution of model. Red grid cells indicate areas where *P. striiformis* was modelled as present recent climate conditions. Grey and white regions indicate where *P. striiformis* has, or has not, been reported, respectively (Pasiecznik et al., 2005; Bebber et al., 2013). (b, d) Conversion of model output to regional scale. For a particular pathogen, white indicates regions modelled as not climatically suitable (Absent) and where the pathogen has not been reported (Absent) (True negative, Specificity), green indicates regions modelled as climatically suitable (Present) and where the pathogen has been reported (Present) (True positive, Sensitivity), yellow indicates regions modelled as climatically suitable (Present) but where the pathogen has not been reported (Absent) (False positive, Type 1 error), and pink indicates regions modelled as not climatically suitable (Absent) but where the pathogen has been reported (Present) (False negative, Type 2 error). A pathogen was modelled as 'present' in a region if it was modelled as 'present' in any grid cell ($j$), for any month ($i$), in the region. (a, b) Temperature-only model. (c, d) Temperature+host model. Proportion True negative, True positive, False positive and False negative for (e) temperature-only and (f) temperature+host outputs. Proportions calculated as the number of regions of a particular colour, relative to the total number of regions per map (396), for each pathogen ($N$ = 67). (g) Comparison of temperature-only and temperature+host output. Decrease (-) in type 1 error refers to regions that transition from yellow to white and increase (+) in type 2 error refers to regions that transition from green to pink, as a result of host restriction (temperature+host). For temperature+host, (h) median gross domestic product based on purchasing power parity (GDP (PPP)) was lower for false positive (yellow) regions than true positive (green) regions (mean difference -0.27 ± 0.037 SE, $t$ = -7.35, df = 87.0, $P$ < 1.03e-10); (i) however, median research output was not (mean difference 0.02 ± 0.100 SE, $t$ = 0.25, df = 123.5, $P$ = 0.81). Boxplot boundaries reflect the inter-quartile range, the horizontal bar is the median. Pathogens restricted by host distributions extracted from EarthStat (Fig S8).

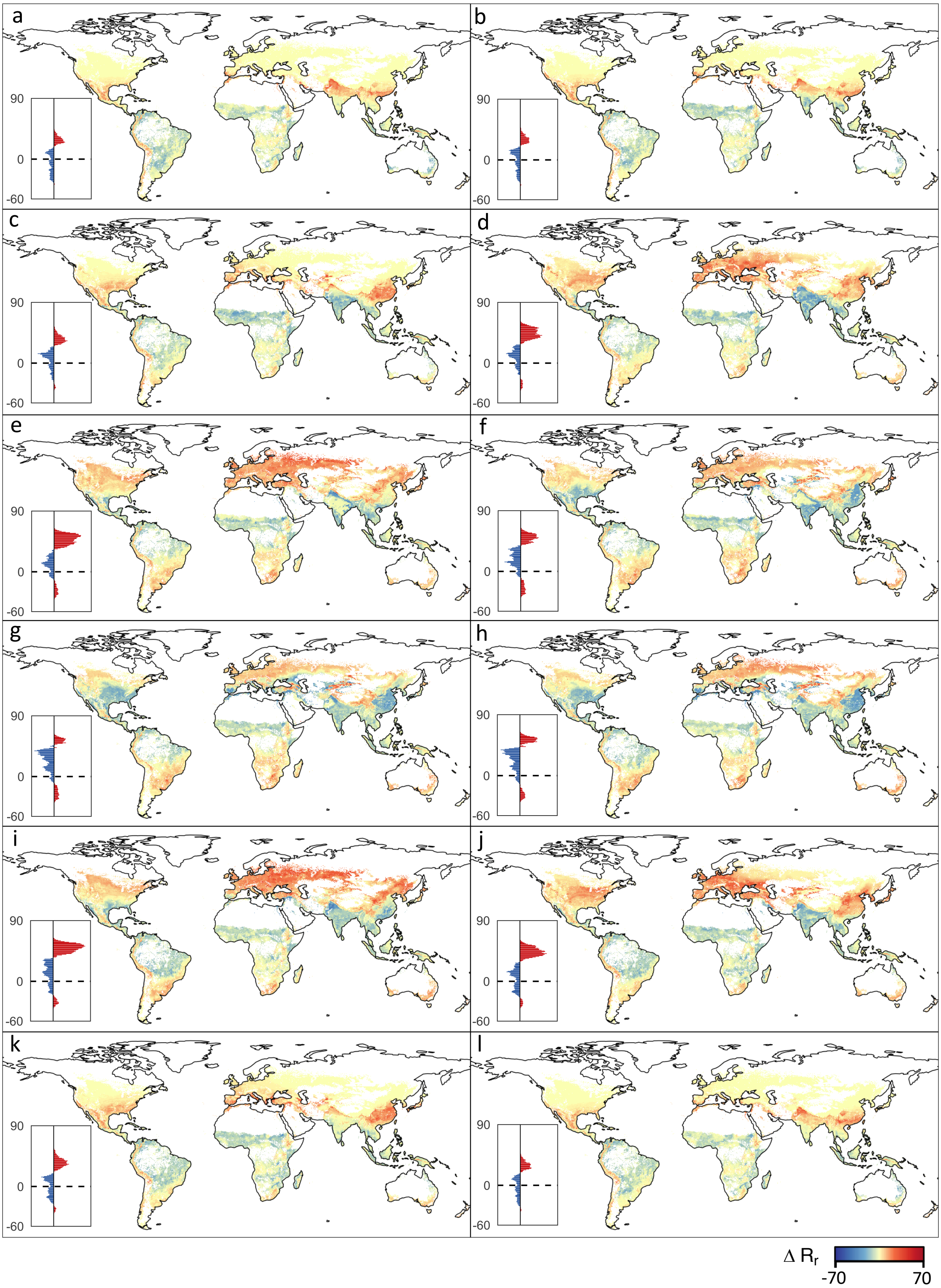

**Fig. S11. Monthly change in global pathogen $R_r$ under RCP 6.0**. (a) Jan., (b) Feb., (c) Mar., (d) Apr., (e) May, (f) June, (g) July, (h) Aug., (i) Sept., (j) Oct., (k) Nov., (l) Dec. $R_r$ defined as the number of pathogens with relative temperature-dependent infection rates ≥ 0.5, i.e. those pathogens with high predicted infection rates. White grid cells contain no hosts (EarthStat, Fig. S8), and were excluded from the analysis. Values on histograms refer to latitude. Red and blue bars reflect increases and decreases in $R_r$, respectively.

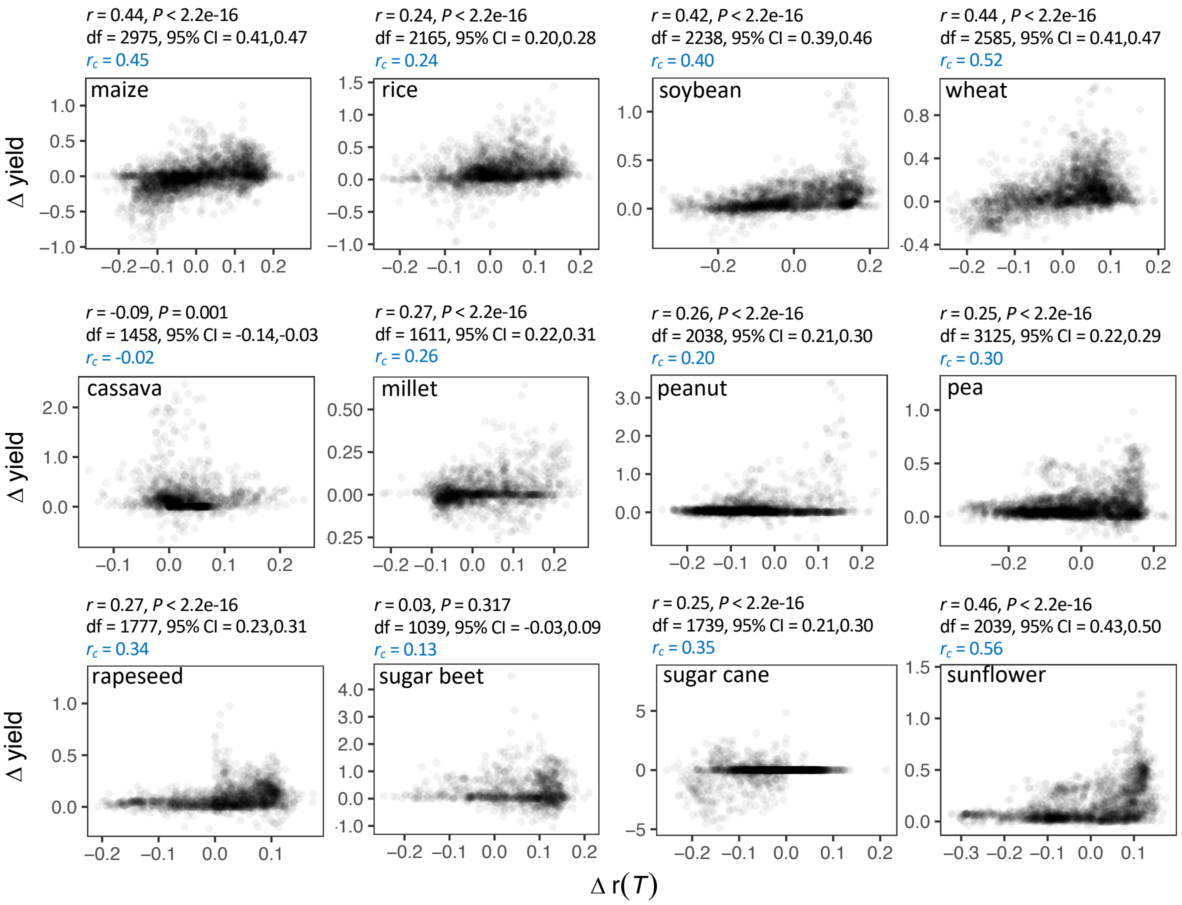

**Fig S12. Pearson correlation (*r*) and spatial cross-correlation ($r_c$) between future crop yield and future temperature-dependent infection risk r(*T*) under RCP 6.0**. Major crops shown in the first row.

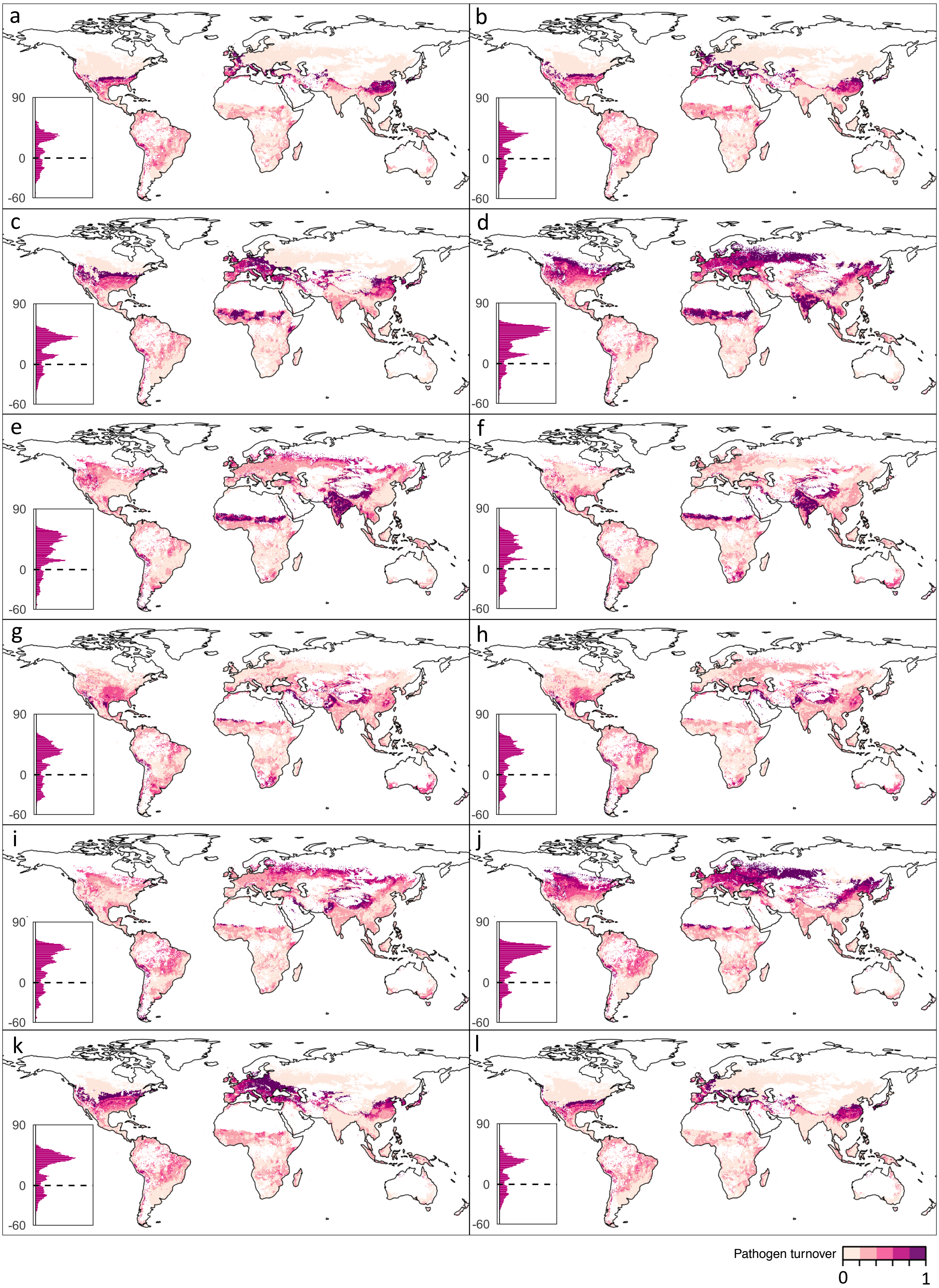

**Fig. S13. Monthly change in global pathogen turnover under RCP 6.0**. (a) Jan., (b) Feb., (c) Mar., (d) Apr., (e) May, (f) June, (g) July, (h) Aug., (i) Sept., (j) Oct., (k) Nov., (l) Dec. Pathogen turnover defined by a modified Jaccard (*J*) index (1 - *J*) of community dissimilarity (see Equation S2). White grid cells contain no hosts (EarthStat, Fig. S8), and were excluded from the analysis. Values on histograms refer to latitude. Pink bars reflect increases in pathogen turnover.

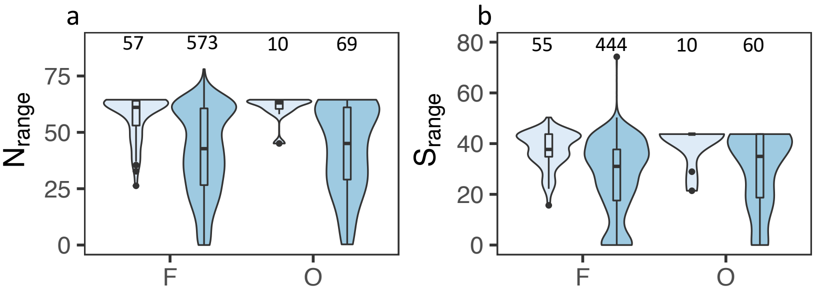

**Fig S14. Violin plots of total (a) northern ($N_{range}$) and (b) southern ($S_{range}$) latitudinal ranges of fungi (F) and oomycetes (O) included in this study**. Light blue represents 67 of the 80 pathogens included in this study. Dark blue represents all fungi and oomycete pathogens included in the CABI Plantwise distribution database. Sample sizes shown above each plot.

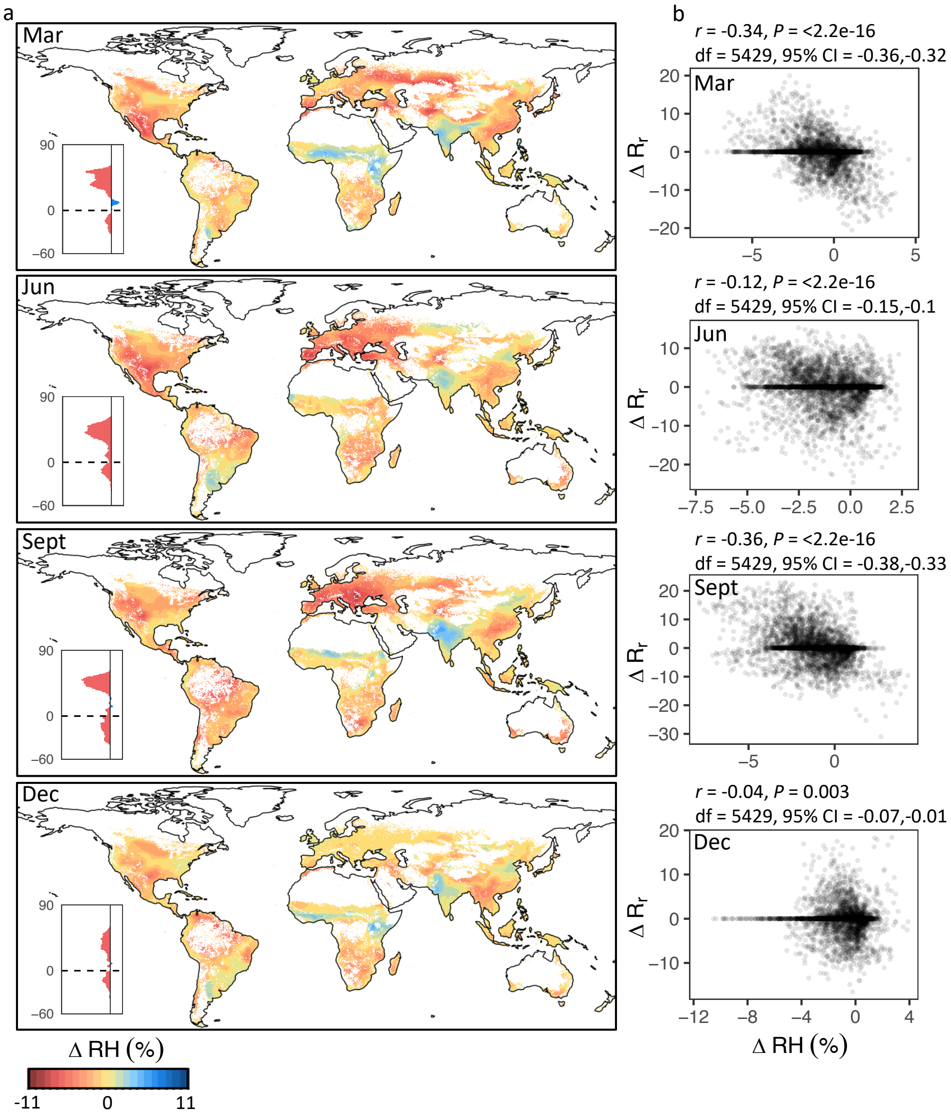

**Fig. S15. Projected monthly change in global near surface RH (1985 - 2070)**. (a) Global distributions. White grid cells contain no hosts (EarthStat, Fig. S8), and were excluded from the analysis. Values on histograms refer to latitude. Blue and pink bars reflect increases and decreases in RH, respectively. (b) Comparison between change in near surface RH and change in $R_r$. Pearson correlations ($r$), df, 95% confidence intervals, and $P$ values shown above figure panels. Grid cells aggregated to 2° spatial resolution to calculate $r$. For further details, see Table S8. Data aggregated to 1° resolution for histograms.

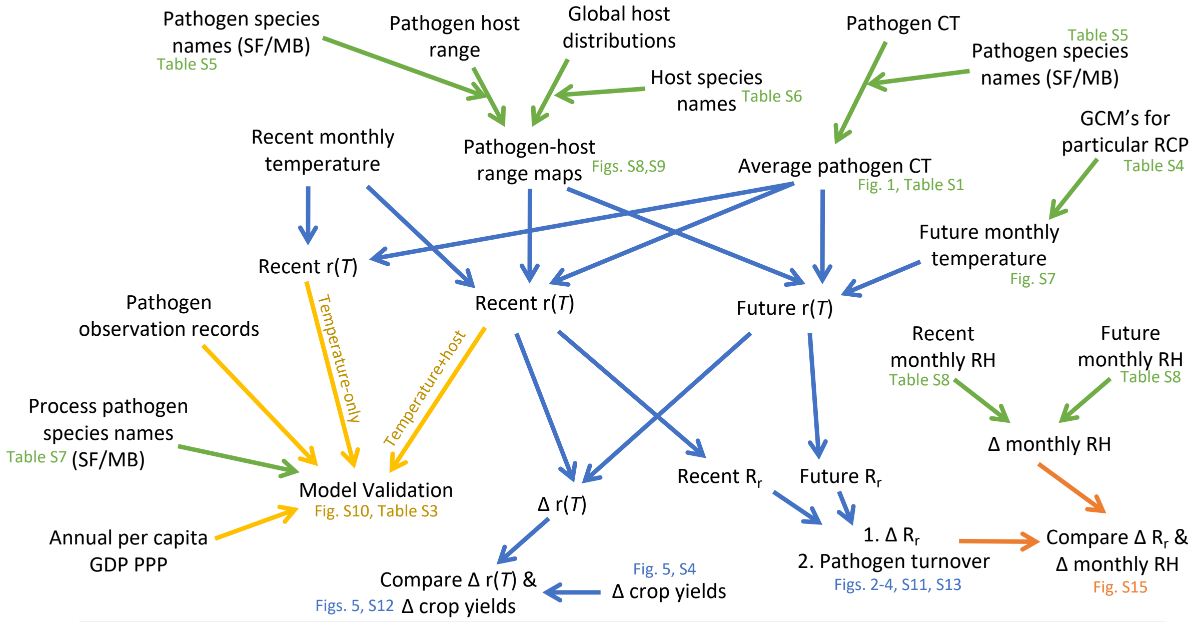

Data Collection

- Crop yields (ISIMIP)
- Crop production (FAO)
- Pathogen CT ($T_{min}$, $T_{opt}$, $T_{max}$)
  - Chaloner *et al.*, (2020)
  - Magarey *et al*., (2005)
  - Viswanath *et al.,* (2017)
  - Boixel *et al*., (2019)
- Pathogen observation records (CABI)
- Recent and future global air surface temperatures (WorldClim)
- Current and future global near surface relative humidity (Climate Data Store)
- Annual per capita GDP at PPP (WorldBank)
- Pathogen host range
  - CABI
  - Magarey *et al*., (2005)
- Global host distributions
  - EarthStat (exc. growing season)
  - MIRCA2000 (inc. growing season)

SF = Species Fungorum/Index Fungorum MB = Mycobank CT = cardinal temperature RH = relative humidity
r(*T*) = relative temperature-dependent infection rate GDP PPP = gross domestic product based on purchasing power parity
$R_r$ = number of pathogens with high temperature-dependent infection risk (r(*T*) ≥ 0.5)

**Fig. S16. Overview of the workflow to prepare pathogen, climate, and host data (green arrows), perform the temperature-dependent infection risk models (blue arrows), perform model validation (yellow arrows), and consider RH on model output (orange arrows)**.

**Table S1. Summary information**. ID refers to pathogen ID in Fig 1. Where multiple data were available, mean $T_{min}$ and/or $T_{opt}$ and/or $T_{max}$ were calculated. Hosts refers to the number of hosts in EarthStat (ES) or MIRCA2000 (MI) database that a pathogen was reported to infect in the Plantwise database.

| **Pathogen** | **Ref** | **ID** | **process** | $T_{min}$ | $T_{opt}$ | $T_{max}$ | **Hosts (ES)** | **Hosts (MI)** |
|---|---|---|---|---|---|---|---|---|
| *Alternaria brassicae* | 16,40 | 19 | infection | 3.80 | 25.00 | 35.23 | 28 | 4 |
| *Alternaria cucumerina* | 16 | 62 | infection | 12.00 | 19.00 | 25.00 | 2 | 0 |
| *Alternaria longipes* | 40 | 16 | infection | 16.50 | 25.75 | 27.00 | 4 | 1 |
| *Alternaria mali* | 16 | 30 | infection | 1.00 | 23.00 | 35.00 | 1 | 0 |
| *Alternaria porri* | 16 | 31 | infection | 1.00 | 23.00 | 35.00 | 3 | 0 |
| *Alternaria radicina* | 40 | 5 | infection | -0.60 | 28.00 | 34.00 | 3 | 1 |
| *Alternaria solani* | 40 | 59 | infection | 5.00 | 19.88 | 25.00 | 6 | 2 |
| *Aphanomyces euteiches* | 40 | 35 | infection | 10.00 | 22.50 | 30.00 | 10 | 0 |
| *Austropuccinia psidii* | 16 | 43 | infection | 1.00 | 21.50 | 30.00 | 2 | 0 |
| *Berkeleyomyces basicola* | 40 | 47 | infection | 15.00 | 20.43 | 29.58 | 44 | 5 |
| *Bipolaris oryzae* | 16,40 | 12 | infection | 15.20 | 26.94 | 37.17 | 2 | 2 |
| *Bipolaris sorokiniana* | 40 | 3 | infection | 9.00 | 29.00 | 34.75 | 33 | 10 |
| *Blumeriella jaapii* | 16,40 | 63 | infection | 6.75 | 19.00 | 29.00 | 4 | 0 |
| *Botryosphaeria dothidea* | 16 | 6 | infection | 8.00 | 28.00 | 35.00 | 8 | 1 |
| *Botryotinia squamosa* | 16,40 | 72 | infection | 2.50 | 17.25 | 25.50 | 2 | 0 |
| *Botrytis cinerea* | 16,40 | 51 | infection | 3.82 | 20.04 | 28.37 | 2 | 1 |
| *Bremia lactucae* | 16 | 78 | infection | 1.00 | 15.00 | 25.00 | 1 | 0 |
| *Ceratocystis fimbriata* | 40 | 20 | infection | 9.75 | 25.00 | 35.13 | 18 | 3 |
| *Cercospora arachidicola* | 16 | 24 | infection | 13.30 | 24.00 | 35.00 | 1 | 1 |
| *Cercospora carotae* | 16 | 25 | infection | 11.00 | 24.00 | 32.00 | 2 | 0 |
| *Colletotrichum acutatum* | 16 | 8 | infection | 7.00 | 27.50 | 35.00 | 16 | 1 |
| *Colletotrichum lindemuthianum* | 40 | 21 | infection | 9.63 | 25.00 | 28.80 | 11 | 1 |
| *Colletotrichum orbiculare* | 16,40 | 26 | infection | 11.50 | 24.00 | 30.00 | 5 | 0 |
| *Didymella arachidicola* | 16 | 65 | infection | 13.30 | 18.50 | 35.00 | 1 | 1 |
| *Didymella pinodes* | 16 | 53 | infection | 1.40 | 20.00 | 35.00 | 6 | 0 |
| *Diplocarpon earlianum* | 16 | 36 | infection | 2.90 | 22.50 | 35.00 | 1 | 0 |
| *Fulvia fulva* | 40 | 34 | infection | 7.20 | 22.53 | 26.70 | 1 | 0 |
| *Fusarium graminearum* | 40 | 46 | infection | 10.42 | 20.91 | 31.32 | 30 | 10 |
| *Fusarium oxysporum f.sp. conglutinans* | 40 | 13 | infection | 17.83 | 26.50 | 34.00 | 2 | 0 |
| *Fusarium oxysporum f.sp. lini* | 40 | 54 | infection | 12.24 | 20.00 | 38.00 | 2 | 0 |
| *Fusarium oxysporum f.sp. lycopersici* | 40 | 4 | infection | 21.00 | 28.19 | 33.00 | 1 | 0 |
| *Fusarium oxysporum f.sp. niveum* | 40 | 29 | infection | 17.30 | 23.38 | 30.00 | 1 | 0 |
| *Fusarium oxysporum f.sp. vasinfectum* | 40 | 2 | infection | 14.20 | 29.30 | 31.60 | 7 | 0 |
| *Fusarium roseum* | 40 | 76 | infection | 0.00 | 15.63 | 30.75 | 15 | 4 |
| *Globisporangium debaryanum* | 40 | 14 | infection | 9.67 | 26.17 | 31.50 | 29 | 6 |
| *Globisporangium ultimum* | 40 | 71 | infection | 2.00 | 17.38 | 32.25 | 34 | 8 |
| *Gymnosporangium juniperi-virginianae* | 16,40 | 50 | infection | 5.50 | 20.20 | 35.00 | 1 | 0 |
| *Leptosphaeria maculans* | 16 | 66 | infection | 2.60 | 18.50 | 35.00 | 6 | 1 |
| *Macrophomina phaseolina* | 40 | 1 | infection | 18.78 | 34.67 | 37.00 | 67 | 13 |
| *Magnaporthe oryzae* | 39 | 9 | lesion development | 7.90 | 27.40 | 34.10 | 3 | 2 |
| *Melampsora lini* | 40 | 64 | infection | 7.00 | 19.00 | 30.00 | 2 | 0 |
| *Monilinia fructicola* | 16 | 55 | infection | 10.00 | 20.00 | 35.00 | 8 | 0 |
| *Mycosphaerella rabiei* | 16 | 22 | infection | 1.00 | 25.00 | 35.00 | 7 | 0 |
| *Nothopassalora personata* | 16 | 56 | infection | 8.00 | 20.00 | 35.00 | 1 | 1 |
| *Peyronellaea obtusa* | 16 | 15 | infection | 1.00 | 26.00 | 35.00 | 8 | 1 |
| *Phakopsora pachyrhizi* | 16 | 32 | infection | 10.00 | 23.00 | 28.00 | 7 | 1 |
| *Phyllosticta ampelicida* | 16 | 10 | infection | 7.00 | 27.00 | 35.00 | 1 | 1 |
| *Phytophthora cactorum* | 16,40 | 37 | infection | 5.50 | 22.50 | 33.17 | 10 | 1 |
| *Phytophthora infestans* | 16,40 | 67 | infection | 3.62 | 18.20 | 31.50 | 5 | 1 |
| *Phytophthora nicotianae* | 40 | 7 | infection | 20.00 | 28.00 | 34.00 | 13 | 0 |
| *Plasmopara viticola* | 16,40 | 41 | infection | 7.88 | 21.83 | 29.33 | 1 | 1 |
| *Pseudoperonospora cubensis* | 16,40 | 60 | infection | 6.30 | 19.75 | 29.00 | 6 | 0 |
| *Puccinia arachidis* | 16 | 17 | infection | 5.00 | 25.00 | 35.00 | 1 | 1 |
| *Puccinia graminis* | 40 | 44 | infection | 13.31 | 21.03 | 27.08 | 5 | 3 |
| *Puccinia hordei* | 40 | 69 | infection | 8.00 | 17.88 | 27.17 | 1 | 1 |
| *Puccinia menthae* | 16 | 79 | infection | 5.00 | 15.00 | 35.00 | 1 | 0 |
| *Puccinia recondita* | 16,40 | 42 | infection | 8.80 | 21.75 | 29.67 | 5 | 3 |
| *Puccinia sorghi* | 40 | 68 | infection | 8.00 | 18.00 | 32.00 | 4 | 1 |
| *Puccinia striiformis* | 16,40 | 80 | infection | 2.60 | 10.50 | 20.78 | 3 | 3 |
| *Pyrenopeziza brassicae* | 16 | 73 | infection | 2.60 | 16.00 | 24.00 | 2 | 1 |
| *Pyrenophora teres* | 16,40 | 39 | infection | 2.60 | 22.00 | 35.00 | 7 | 3 |
| *Pythium arrhenomanes* | 40 | 74 | infection | 12.00 | 16.00 | 38.00 | 17 | 7 |
| *Rhizoctonia solani* | 40 | 11 | infection | 21.07 | 26.94 | 34.10 | 40 | 13 |

| | | | | | | | | |
|---|---|---|---|---|---|---|---|---|
| *Rhizopus stolonifer* | 40 | 49 | infection | 7.52 | 20.29 | 27.23 | 31 | 5 |
| *Rhynchosporium secalis* | 16 | 38 | infection | 2.60 | 22.50 | 30.00 | 4 | 3 |
| *Sclerospora graminicola* | 40 | 61 | infection | 11.00 | 19.25 | 34.00 | 6 | 3 |
| *Sclerotinia sclerotiorum* | 16,40 | 40 | infection | 0.38 | 22.00 | 28.63 | 63 | 7 |
| *Septoria glycines* | 16 | 23 | infection | 10.00 | 25.00 | 35.00 | 1 | 1 |
| *Sporisorium cruentum* | 40 | 33 | infection | 15.00 | 23.00 | 35.00 | 3 | 2 |
| *Sporisorium reilianum* | 40 | 27 | infection | 16.00 | 24.00 | 36.00 | 6 | 2 |
| *Sporisorium sorghi* | 40 | 28 | infection | 12.50 | 23.50 | 35.00 | 2 | 1 |
| *Stigmina carpophila* | 16 | 18 | infection | 5.00 | 25.00 | 35.00 | 4 | 0 |
| *Synchytrium endobioticum* | 40 | 77 | infection | 9.60 | 15.60 | 26.18 | 1 | 1 |
| *Urocystis cepulae* | 40 | 70 | infection | 10.00 | 17.50 | 27.20 | 3 | 0 |
| *Uromyces viciae-fabae* | 40 | 57 | infection | 4.00 | 20.00 | 33.00 | 5 | 0 |
| *Ustilago avenae* | 40 | 52 | infection | 5.29 | 20.04 | 30.33 | 2 | 1 |
| *Venturia inaequalis* | 16,40 | 58 | infection | 3.50 | 19.88 | 29.00 | 2 | 0 |
| *Venturia pyrina* | 16 | 45 | infection | 1.00 | 21.00 | 35.00 | 2 | 0 |
| *Wilsoniana occidentalis* | 16 | 75 | infection | 6.00 | 16.00 | 28.00 | 1 | 0 |
| *Zymoseptoria tritici* | 42 | 48 | growth in culture | 4.90 | 20.30 | 33.50 | 2 | 2 |

**Table S2. The number of pathogens included for each host in analysis of change in infection rate under RCP 6.0**.

| Host | Pathogens |
|---|---|
| Cassava | 3 |
| Peanut | 10 |
| Maize | 13 |
| Millet | 4 |
| Pea | 12 |
| Rapeseed | 7 |
| Rice | 8 |
| Soybean | 10 |
| Sugar beet | 7 |
| Sugar cane | 8 |
| Sunflower | 8 |
| Wheat | 14 |

**Table S3. Pathogen distribution model accuracy**. Median and interquartile range for pathogen projected global distributions, for distribution based on temperature only, or temperature and host availability.

| Classification | Temperature only model | Temperature+host model |
|---|---|---|
| True positive (Sensitivity) | 0.23 (0.16 – 0.30) | 0.21 (0.15 – 0.28) |
| True negative (Specificity) | 0.04 (0.03 – 0.09) | 0.27 (0.18 – 0.35) |
| False positive (Type 1 error) | 0.69 (0.63 – 0.77) | 0.47 (0.37 – 0.57) |
| False negative (Type 2 error) | 0.00 (0.00 – 0.00) | 0.01 (0.01-0.03) |

**Table S4. All global climate models (GCMs) of Representative Concentration Pathways (RCP) 2.6, 4.5, 6.0 and 8.5 used in this study**. 'Y' detonates GCMs that were available for each RCP. For each GCM, average future monthly temperature was calculated as the mid-point of average maximum and minimum monthly temperature, as no average estimates were available.

| GCM | RCP 2.6 | RCP 4.5 | RCP 6.0 | RCP 8.5 |
|---|---|---|---|---|
| ACCESS1-0 | | Y | | Y |
| BCC-CSM1-1 | Y | Y | Y | Y |
| CCSM4 | Y | Y | Y | Y |
| CESM1-CAM5-1-FV2 | | Y | | |
| CNRM-CM5 | Y | Y | | Y |
| GFDL-CM3 | Y | Y | | Y |
| GFDL-ESM2G | Y | Y | Y | |
| GISS-E2-R | Y | Y | Y | Y |
| HadGEM2-AO | Y | Y | Y | Y |
| HadGEM2-CC | | Y | | Y |
| HadGEM2-ES | Y | Y | Y | Y |
| INMCM4 | | Y | | Y |
| IPSL-CM5A-LR | Y | Y | Y | Y |
| MIROC-ESM-CHEM | Y | Y | Y | Y |
| MIROC-ESM | Y | Y | Y | Y |
| MIROC5 | Y | Y | Y | Y |
| MPI-ESM-LR | Y | Y | | Y |
| MRI-CGCM3 | Y | Y | Y | Y |
| NorESM1-M | Y | Y | Y | Y |

**Table S5. Updated names of pathogens recorded in Magarey et al. (2005)**. Species Fungorum (SF) was used to update species names to ensure correcting matching to species in Magarey et al. (2005). Where species were not reported in SF, MycoBank (MB) was also used as an alternative. Species discovery name author(s) and sanction name author(s) are not reported in Magarey et al., (2005) and so were not considered here.

| Species name from Magarey et al. (2005) | Updated species name | Source |
|---|---|---|
| *Puccinia psidii* | *Austropuccinia psidii* | SF |
| *Coccomyces hiemalis* | *Blumeriella jaapii* | SF |
| *Botrytis squamosa* | *Botryotinia squamosa* | SF |
| *Mycosphaerella pinodes* | *Didymella pinodes* | SF |
| *Ascochyta rabiei* | *Mycosphaerella rabiei* | SF |
| *Cercosporidium personatum* | *Nothopassalora personata* | SF |
| *Botryosphaeria obtusa* | *Peyronellaea obtuse* | SF |
| *Guignardia bidwellii* | *Phyllosticta ampelicida* | SF |
| *Wilsonomyces carpophilus* | *Stigmina carpophila* | SF |
| *Venturia pirina* | *Venturia pyrina* | MB |
| *Albugo occidentalis* | *Wilsoniana occidentalis* | SF |

**Table S6. Hosts and assigned species names**. All hosts below were extracted in EarthStat, and a subset of hosts were extracted from MIRCA2000. Host names specified in EarthStat and MIRCA2000 were assigned species names below to enable matching to the Plantwise database, when determining plant-pathogen interactions. Hosts listed below were only included in climate models if they were recorded to be a host of at least one pathogen included in this study. Some hosts present in EarthStat and MIRCA2000 were excluded from the table below, see Methods for further details.

| EarthStat (MIRCA2000) host name | Assigned species name(s) |
|---|---|
| abaca | *Musa textilis* |
| agava | *Agave americana, Agave cantala, Agave foetida, Agave fourcroydes, Agave lechuguilla, Agave letonae* |
| alfalfa | *Medicago sativa* |
| almond | *Amygdalus communis, Prunus amygdalus, Prunus communis, Prunus dulcis* |
| apple | *Malus communis, Malus pumila, Malus sylvestris, Pyrus malus, Malus domestica* |
| apricot | *Prunus armeniaca* |
| areca | *Areca catechu* |
| artichoke | *Cynara scolymus* |
| asparagus | *Asparagus officinalis* |
| avocado | *Persea americana* |
| bambara | *Vigna subterranea, Voandzeia subterranea* |
| banana | *Musa sapientum, Musa paradisiaca, Musa cavendishii, Musa acuminata, Musa nana* |
| barley (barley) | *Hordeum disticum, Hordeum distichum, Hordeum hexasticum, Hordeum vulgare* |
| bean | *Phaseolus vulgaris, Phaseolus lunatus, Phaseolus angularis, Vigna angularis, Phaseolus aureus, Phaseolus mungo, Vigna mungo, Phaseolus coccineus, Phaseolus calcaratus, Vigna umbellata, Phaseolus aconitifolius, Phaseolus acutifolius, Vigna radiata, Vigna aconitifolia* |
| beet forage | *Beta vulgaris* (excluding varieties) |
| blueberry | *Vaccinium corymbosum, Vaccinium myrtillus* |
| brazil nut | *Bertholletia excelsa* |
| broadbean | *Vicia faba, Vicia faba* var. *equina, Vicia faba* var. *major, Vicia faba* var. *minor* |
| buckwheat | *Fagopyrum esculentum* |
| cabbage | *Brassica chinensis, Brassica oleracea* var. *alboglabra, Brassica oleracea* var. *capitata, Brassica oleracea* var. *gemmifera, Brassica oleracea* var. *gongylodes, Brassica oleracea* var. *italica, Brassica oleracea* var. *viridis* |
| cabbage, forage | *Brassica chinensis, Brassica oleracea* (excluding varieties) |
| canary seed | *Phalaris canariensis* |
| carob | *Ceratonia siliqua* |
| carrot | *Daucus carota* |
| carrot, forage | *Daucus carota* |
| cashew | *Anacardium occidentale* |
| cashewapple | *Anacardium occidentale* |
| cassava (cassava) | *Manihot esculenta, Manihot dulcis, Manihot palmata, Manihot utilissima* |
| castor | *Ricinus communis* |
| cauliflower | *Brassica oleracea* var. *botrytis* |
| cherry | *Prunus avium, Cerasus avium* var. *duracina, Cerasus avium* var. *juliana* |
| chestnut | *Castanea vesca, Castanea vulgaris, Castanea sativa* |
| chickpea | *Cicer arietinum* |
| chicory | *Cichorium intybus, Cichorium sativum* |
| chille etc | *Capsicum annuum, Capsicum fructescens, Capsicum frutescens, Pimenta officinalis, Pimenta dioica* |
| cinnamon | *Cinnamomum zeylanicum, Cinnamomum cassia* |
| citrus nes | *Citrus bergamia, Citrus medica* var. *cedrata, Citrus medica, Citrus myrtifolia, Citrus aurantium, Fortunella japonica* |
| clove | *Eugenia caryophyllata, Caryophyllus aromaticus, Syzygium aromaticum* |
| clover | *Trifolium* |
| cocoa (cocoa) | *Theobroma cacao* |
| coconut | *Cocos nucifera* |
| coffee (coffee) | *Coffea arabica, Coffea liberica, Coffea robusta* |
| coir | *Cocos nucifera* |
| cotton (cotton) | *Gossypium hirsutum, Gossypium barbadense, Gossypium arboreum, Gossypium herbaceum* |
| cowpea | *Vigna sinensis, Dolichos sinensis, Vigna unguiculata* |
| cranberry | *Vaccinium macrocarpon, Vaccinium oxycoccus* |
| cucumber etc | *Cucumis sativus* |
| currant | *Ribes nigrum, Ribes rubrum* |
| date (date palm) | *Phoenix dactylifera* |
| eggplant | *Solanum melongena* |
| fig | *Ficus carica* |
| flax | *Linum usitatissimum* |
| fonio | *Digitaria exilis, Digitaria iburua* |
| garlic | *Allium sativum* |
| ginger | *Zingiber officinale* |

| | |
|---|---|
| gooseberry | *Ribes grossularia, Ribes uva-crispa* |
| grape (grape/vine) | *Vitis vinifera* |
| grapefruit etc | *Citrus grandis, Citrus maxima, Citrus paradisi* |
| green bean | *Phaseolus acutifolius, Phaseolus coccineus, Phaseolus lunatus, Phaseolus vulgaris, Vigna aconitifolia, Vigna angularis, Vigna mungo, Vigna radiata, Vigna subterranea, Vigna umbellata, Vigna unguiculata, Vigna unguiculata* subsp. *sesquipedalis, Vigna vexillata* |
| green broadbean | *Vicia faba* |
| green corn | *Zea mays, Zea mays* subsp. *mays* |
| green onion | *Allium cepa, Allium cepa* var. *aggregatum, Allium ascalonicum, Allium fistulosum* |
| green pea | *Pisum sativum* |
| groundnut (groundnut/peanut) | *Arachis hypogaea* |
| hazelnut | *Corylus avellana* |
| hemp | *Cannabis sativa* |
| hemp seed | *Cannabis sativa* |
| hop | *Humulus lupulus* |
| jute | *Corchorus capsularis, Corchorus olitorius* |
| kapok fiber | *Ceiba pentandra* |
| kapok seed | *Ceiba pentandra* |
| karite | *Butyrospermum parkii, Vitellaria paradoxa* |
| kiwi | *Actinidia chinensis* |
| kolanut | *Cola acuminata, Cola nitida, Cola vera* |
| lemon, lime | *Citrus limon, Citrus aurantifolia, Citrus aurantiifolia, Citrus limetta, Citrus medica* |
| lentil | *Lens esculenta, Lens culinaris, Ervum lens, Lens culinaris* subsp. *culinaris* |
| lettuce | *Lactuca sativa, Cichorium intybus* var. *foliosum, Cichorium intybus, Cichorium endivia* var. *crispa, Cichorium endivia* var. *latifolia, Cichorium endivia* |
| linseed | *Linum usitatissimum* |
| lupin | *Lupinus albus, Lupinus angustifolius, Lupinus arboreus, Lupinus luteus, Lupinus polyphyllus* |
| maize (maize) | *Zea mays, Zea mays* subsp. *mays* |
| maize, forage | *Zea mays, Zea mays* subsp. *mays* |
| mango | *Mangifera indica* |
| mate | *Ilex paraguariensis* |
| melon etc | *Cucumis melo* |
| melon seed | *Cucumis melo* |
| millet (millet) | *Eleusine coracana, Panicum miliaceum, Pennisetum glaucum, Setaria italica, Echinocloa frumentacea, Echinochloa frumentacea, Paspalum scrobiculatum* |
| mustard | *Brassica alba, Brassica hirta, Sinapis alba, Brassica nigra, Sinapis nigra* |
| nutmeg | *Myristica fragrans, Elettaria cardamomum, Aframomum angustifolium, Aframomum hambury, Amomum aromaticum, Amomum cardamomum, Aframomum melegueta* |
| oats | *Avena sativa* |
| oil palm (oil palm) | *Elaeis guineensis* |
| okra | *Abelmoschus esculentus, Hibiscus esculentus* |
| olive | *Olea europaea, Olea europaea* subsp. *europaea* |
| onion | *Allium cepa, Allium cepa* var. *aggregatum* |
| orange | *Citrus sinensis, Citrus aurantium* |
| papaya | *Carica papaya* |
| pea (pulses) | *Pisum sativum, Pisum arvense* |
| peach etc | *Prunus persica, Amygdalus persica, Persica laevis* |
| pear | *Pyrus communis* |
| pepper | *Piper longum, Piper nigrum* |
| peppermint | *Mentha piperita* |
| persimmon | *Diospyros kaki, Diospyros virginiana* |
| pigeon pea | *Cajanus cajan* |
| pimento | *Capsicum annuum, Capsicum frutescens, Pimenta officinalis, Pimenta dioica* |
| pineapple | *Ananas comosus, Ananas sativus* |
| pistachio | *Pistacia vera* |
| plantain | *Musa paradisiaca* |
| plum | *Prunus domestica, Prunus spinosa* |
| popcorn | *Zea mays* var. *everta, Zea mays, Zea mays* subsp. *mays* |
| poppy | *Papaver somniferum* |
| potato (potato) | *Solanum tuberosum* |
| pumpkin etc | *Cucurbita pepo, Cucurbita maxima, Cucurbita moschata* |
| pyrethrum | *Chrysanthemum cinerariifolium, Tanacetum cinerariifolium* |
| quince | *Cydonia oblonga, Cydonia vulgaris, Cydonia japonica, Chaenomeles japonica* |
| quinoa | *Chenopodium quinoa* |
| ramie | *Boehmeria nivea, Boehmeria tenacissima* |
| rapeseed (rapeseed/canola) | *Brassica napus* var. *oleifera, Brassica napus* var. *napus* |
| raspberry | *Rubus idaeus* |
| rice (rice) | *Oryza sativa* |
| rubber | *Hevea brasiliensis* |
| rye (rye) | *Secale cereale* |

| | |
|---|---|
| rye, forage | *Lolium multiflorum, Lolium perenne* |
| safflower | *Carthamus tinctorius* |
| sesame | *Sesamum indicum* |
| sisal | *Agave sisalana* |
| sorghum (sorghum) | *Sorghum guineense, Sorghum vulgare, Sorghum dura, Sorghum bicolor* |
| sorghum, forage | *Sorghum guineense, Sorghum vulgare, Sorghum dura, Sorghum bicolor* |
| sour cherry | *Prunus cerasus, Cerasus acida* |
| soybean (soybean) | *Glycine soja, Glycine max* |
| spinach | *Spinacia oleracea* |
| strawberry | *Fragaria ananassa, Fragaria chiloensis, Fragaria moschata, Fragaria vesca, Fragaria virginiana* |
| stringbean | *Phaseolus vulgaris* |
| sugar beet (sugar beet) | *Beta vulgaris* var. *altissima, Beta vulgaris* var. *saccharifera* |
| sugarcane (sugarcane) | *Saccharum officinarum* |
| sunflower (sunflower) | *Helianthus annuus* |
| swede, forage | *Brassica napus* var. *napobrassica* |
| sweet potato | *Ipomoea batatas* |
| tang etc | *Citrus reticulata, Citrus unshiu* |
| taro | *Colocasia esculenta* |
| tea | *Camellia sinensis, Thea sinensis, Thea assaamica* |
| tobacco | *Nicotiana tabacum* |
| tomato | *Lycopersicon esculentum, Solanum lycopersicum* |
| tung | *Aleurites cordata, Aleurites fordii, Vernicia fordii* |
| turnip, forage | *Brassica rapa* var. *rapifera, Brassica rapa* |
| vanilla | *Vanilla planifolia, Vanilla pompona* |
| vetch | *Vicia sativa* |
| walnut | *Juglans regia* |
| watermelon | *Citrullus vulgaris, Citrullus lanatus* |
| wheat (wheat) | *Triticum aestivum, Triticum durum, Triticum spelta* |
| yam | *Dioscorea rotundata, Dioscorea alata, Dioscorea bulbifera, Dioscorea esculenta, Dioscorea trifida, Dioscorea batatas, Dioscorea cayenensis* |
| yautia | *Xanthosoma sagittifolium* |

**Table S7. Pathogen names updated in the CABI Plantwise database to improve matching to the Pathogen dataset**. *Species excluded from model validation and pathogen sampling bias analyses due to no observational data available.

| Species name in the CABI Plantwise | Updated species name |
|---|---|
| *Botryosphaeria obtusa* | *Peyronellaea obtusa* |
| *Cochliobolus miyabeanus* | *Bipolaris oryzae* |
| *Cochliobolus sativus* | *Bipolaris sorokiniana* |
| *Didymosphaeria arachidicola** | *Didymella arachidicola* |
| *Gibberella pulicaris* | *Fusarium roseum* |
| *Gibberella zeae* | *Fusarium graminearum* |
| *Guignardia bidwellii* | *Phyllosticta ampelicida* |
| *Magnaporthe grisea* | *Magnaporthe oryzae* |
| *Mycosphaerella arachidis* | *Cercospora arachidicola* |
| *Mycosphaerella graminicola* | *Zymoseptoria tritici* |
| *Mycosphaerella pinodes* | *Didymella pinodes* |
| *Passalora fulva* | *Fulvia fulva* |
| *Puccinia psidii* | *Austropuccinia psidii* |
| *Pythium debaryanum* | *Globisporangium debaryanum* |
| *Pythium ultimum* | *Globisporangium ultimum* |
| *Sphacelotheca reiliana* | *Sporisorium reilianum* |
| *Thanatephorus cucumeris* | *Rhizoctonia solani* |
| *Thielaviopsis basicola* | *Berkeleyomyces basicola* |

**Table S8. All recent (R) and future (RCP 6.0) near surface RH model-ensemble combinations used in this study**. Time periods (TP) show yyyymm – yyyymm. TP Estimated calculated as mean of TP Extracted. Change in RH calculated as future RH minus recent RH. *196601-197012; 197101-197512; 197601-198012; 198101-198512; 198601-199012; 199101-199512; 199601-200012; 200101-200512. **195101-197512; 197601-200012. ***195912-198411; 198412-200511. ****195912-198411; 198412-200512.

| Model | Ensemble | TP Downloaded (R) | TP Extracted (R) | TP Estimated (R) | TP Downloaded (RCP 6.0) | TP Extracted (RCP 6.0) | TP Estimated (RCP 6.0) |
|---|---|---|---|---|---|---|---|
| bcc-csm1-1 (BCC, China) | r1i1p1 | 185001-201212 | 197001 - 200012 | 1985 | 200601-209912 | 206001 - 208012 | 2070 |
| bcc-csm1-1m (BCC, China) | r1i1p1 | 185001-201212 | 197001 - 200012 | 1985 | 200601-210012 | 206001 - 208012 | 2070 |
| CCSM4 (NCAR, USA) | r6i1p1 | 185001-200512 | 197001 - 200012 | 1985 | 200601-210012 | 206001 - 208012 | 2070 |
| CESM1-CAM5 (NCAR, USA) | r1i1p1 | 185001-200512 | 197001 - 200012 | 1985 | 200601-210012 | 206001 - 208012 | 2070 |
| CESM1-CAM5 (NCAR, USA) | r2i1p1 | 185001-200512 | 197001 - 200012 | 1985 | 200601-210012 | 206001 - 208012 | 2070 |
| CESM1-CAM5 (NCAR, USA) | r3i1p1 | 185001-200512 | 197001 - 200012 | 1985 | 200601-210012 | 206001 - 208012 | 2070 |
| CSIRO-Mk3-6-0 (CSIRO, Australia) | r1i1p1 | 185001-200512 | 197001 - 200012 | 1985 | 200601-210012 | 206001 - 208012 | 2070 |
| CSIRO-Mk3-6-0 (CSIRO, Australia) | r2i1p1 | 185001-200512 | 197001 - 200012 | 1985 | 200601-210012 | 206001 - 208012 | 2070 |
| CSIRO-Mk3-6-0 (CSIRO, Australia) | r3i1p1 | 185001-200512 | 197001 - 200012 | 1985 | 200601-210012 | 206001 - 208012 | 2070 |
| CSIRO-Mk3-6-0 (CSIRO, Australia) | r4i1p1 | 185001-200512 | 197001 - 200012 | 1985 | 200601-210012 | 206001 - 208012 | 2070 |
| CSIRO-Mk3-6-0 (CSIRO, Australia) | r5i1p1 | 185001-200512 | 197001 - 200012 | 1985 | 200601-210012 | 206001 - 208012 | 2070 |
| CSIRO-Mk3-6-0 (CSIRO, Australia) | r6i1p1 | 185001-200512 | 197001 - 200012 | 1985 | 200601-210012 | 206001 - 208012 | 2070 |
| CSIRO-Mk3-6-0 (CSIRO, Australia) | r7i1p1 | 185001-200512 | 197001 - 200012 | 1985 | 200601-210012 | 206001 - 208012 | 2070 |
| CSIRO-Mk3-6-0 (CSIRO, Australia) | r8i1p1 | 185001-200512 | 197001 - 200012 | 1985 | 200601-210012 | 206001 - 208012 | 2070 |
| CSIRO-Mk3-6-0 (CSIRO, Australia) | r9i1p1 | 185001-200512 | 197001 - 200012 | 1985 | 200601-210012 | 206001 - 208012 | 2070 |
| CSIRO-Mk3-6-0 (CSIRO, Australia) | r10i1p1 | 185001-200512 | 197001 - 200012 | 1985 | 200601-210012 | 206001 - 208012 | 2070 |
| GFDL-ESM2G (NOAA, USA) | r1i1p1 | 196601-200512* | 197001 - 200012 | 1985 | 206601-207012 | 207001-207012 | 2070 |
| GFDL-ESM2M (NOAA, USA) | r1i1p1 | 196601-200512* | 197001 - 200012 | 1985 | 206601-207012 | 207001-207012 | 2070 |
| GISS-E2-H (NASA, USA) | r1i1p1 | 195101-200512 | 197001 - 200012 | 1985 | 205101-210012 | 206001 - 208012 | 2070 |
| GISS-E2-H (NASA, USA) | r1i1p2 | 195101-200512 | 197001 - 200012 | 1985 | 205101-210012 | 206001 - 208012 | 2070 |
| GISS-E2-H (NASA, USA) | r1i1p3 | 195101-200512 | 197001 - 200012 | 1985 | 205101-210012 | 206001 - 208012 | 2070 |
| GISS-E2-R (NASA, USA) | r1i1p1 | 195101-200012** | 197001 - 200012 | 1985 | 205101-207512 | 206501 - 207512 | 2070 |
| GISS-E2-R (NASA, USA) | r1i1p2 | 195101-200012** | 197001 - 200012 | 1985 | 205101-207512 | 206501 - 207512 | 2070 |
| GISS-E2-R (NASA, USA) | r1i1p3 | 195101-200012** | 197001 - 200012 | 1985 | 205101-207512 | 206501 - 207512 | 2070 |
| HadGEM2-ES (UK Met Office, UK) | r1i1p1 | 195912-200511*** | 197001 - 200012 | 1985 | 206112-208611 | 206201 - 207812 | 2070 |

| | | | | | | | |
|---|---|---|---|---|---|---|---|
| HadGEM2-ES (UK Met Office, UK) | r2i1p1 | 195912-200512**** | 197001 - 200012 | 1985 | 205512-208011 | 206101 - 207912 | 2070 |
| HadGEM2-ES (UK Met Office, UK) | r3i1p1 | 195912-200512**** | 197001 - 200012 | 1985 | 205512-208011 | 206101 - 207912 | 2070 |
| HadGEM2-ES (UK Met Office, UK) | r4i1p1 | 195912-200511*** | 197001 - 200012 | 1985 | 205512-208011 | 206101 - 207912 | 2070 |
| IPSL-CM5A-LR (IPSL, France) | r1i1p1 | 185001-200512 | 197001 - 200012 | 1985 | 200601-210012 | 206001 - 208012 | 2070 |
| IPSL-CM5A-MR (IPSL, France) | r1i1p1 | 185001-200512 | 197001 - 200012 | 1985 | 200601-210012 | 206001 - 208012 | 2070 |
| NorESM1-M (NCC, Norway) | r1i1p1 | 185001-200512 | 197001 - 200012 | 1985 | 200601-210012 | 206001 - 208012 | 2070 |
| NorESM1-ME (NCC, Norway) | r1i1p1 | 185001-200512 | 197001 - 200012 | 1985 | 205101-210112 | 206001 - 208012 | 2070 |

### Appendix

#### The influence of canopy moisture on estimates of temperature-dependent infection risk

##### Introduction

In this study, we have employed temperature response functions of plant pathogen infection rates, r($T$), to project potential shifts in infections at the global scale. While temperature is a major determinant of plant disease risk[1], high humidity or leaf wetness are required for spore germination and host penetration by many fungal and oomycete plant pathogens[2]. Therefore, infection risk models that employ temperature alone could give biased estimates of spatiotemporal trends in infection risk, if humidity changes unevenly over the region of interest. The scaling of moisture availability for spore germination and subsequent hyphal growth and infection, from the leaf surface, to the plant canopy, to the crop and the modelled geographical grid cell is a complex topic beyond the scope of the present analysis. We can, however, compare results from infection models employing both temperature and leaf wetness, with those from models employing temperature alone, to investigate how changes in moisture affect estimates of temperature-driven infection rates.

##### Methods

For this model comparison, we employed historical estimates of canopy temperature and canopy surface moisture (leaf wetness). We used historical climate reanalyses because we can be more confident in them than future climate projections. The Japanese 55-year Reanalysis (JRA55) dataset[3], previously used for modelling historical trends in fungal pathogen infection rate[4], provides global historical climate data at 0.5625° spatial resolution and 3-hourly intervals from 00:00h on 1st January 1958 onwards. JRA-55 historical climate re-analysis data was obtained from the Research Data Archive at the National Center for Atmospheric Research (Boulder, Colorado) at https://rda.ucar.edu/.

We obtained canopy temperature (K) and canopy surface moisture ($m^3$ $m^{-2}$, converted to g $m^{-2}$ for analysis) for a 60-year period (00:00h on 1st January 1958 to 21:00h on 31st December 2017) for two geographical regions, in North America and East Africa. The North American region covers the 'wheat belt' stretching from Texas north to North Dakota and into Saskatchewan and Manitoba in Canada (Fig. A1a). The North American region spanned 94.79 – 104.90 °W and 32.01 – 52.23 °N, on the boundary between the drier western and wetter eastern halves of the continent. The dataset comprised 18 model grid cells longitudinally and 36 grid cells latitudinally, with 175,320 timepoints. Winter wheat predominates in the southern US states of Texas, Oklahoma, Kansas, and Nebraska, while spring wheat is more common in the northern states of South and North Dakota, and in Canada[5,6].

The East African region covers the maize-growing area stretching from South Africa to Ethiopia (Fig. A1b). The region spanned 23.91 – 40.21°E and 32.01 °S – 14.04 °N, comprising 29 model grid cells longitudinally and 82 grid cells latitudinally. We chose these regions because they each cover a large latitudinal range (20° in North America, 46° in East Africa), and are dominated by a single major grain crop (wheat and maize, respectively). We did not attempt to model 3-hourly infection rate for 60 years over the entire globe due to the very large sizes of the required databases.

In each region, we modelled r(*T*) by a rust fungus infecting the major grain crop in that area. We selected rusts because infection of crops by rust urediniospores is strongly constrained by moisture: spore germination can only occur in the presence of free water on the leaf surface, and desiccation kills germinated spores [4,7]. Therefore, any prediction biases caused by omission of moisture effects in our global models should be most strongly evident for these pathogens. For North American wheat we modelled stripe (yellow) rust, caused by *Puccinia striiformis*[8]. For East African maize we modelled common rust of maize, caused by *Puccinia sorghi*[9]. The urediniospores of both these fungi require moisture to germinate. The cardinal temperatures for each pathogen are given in Table S1.

We compared three infection models. The first model estimated r(*T*) at each 3-hour timestep using the beta function of cardinal temperatures, but set r(*T*) to zero for any periods when canopy surface moisture was zero. The second model estimated r(*T*) at each 3-hour timestep without canopy surface moisture restrictions, i.e., leaf wetness requirements were ignored. The third model replicated our global model and estimated r(*T*) from mean monthly temperature, without considering canopy moisture. We designate the results of these models $r_{wet}$, $r_{all}$ and $r_{mon}$, respectively. For North American wheat, we modelled only early to late spring (February to May, inclusive), as stripe rust infections are most commonly reported during this period [10]. For East African maize, we modelled only the maize growing season, using estimates of growing season from the MIRCA2000 database. For Ethiopia and Sudan the growing season starts in June and ends in October. For Kenya, Uganda and Tanzania it starts in April and ends in August. For DRC it starts in October and ends in January. For all more southerly countries, including South Africa, the growing season starts in January and ends in April. The models were compared by correlating annual mean r(*T*) per grid cell during the crop growing season, and by correlating the 60-year trend in r(T) per grid cell.

**Results**

In the North America region, mean spring temperature declines with latitude (Fig. A2a), and increases over the analysis period (1958-2017), particularly in the central western sector (Fig. A2b). Mean canopy moisture is greater in far eastern edge of the region (Fig. A2c), and there has been little change over time with an average trend of 0.00 g $m^{-2}$ $y^{-1}$ over the analysis period (Fig. A2d). In the East Africa region, there is no strong latitudinal trend in the mean growing season temperature (Fig. A3a), but an apparent cooling trend in the southern portion and warming trend in the north (Fig. A3b). Mean growing season canopy moisture shows no latitudinal trend (Fig. A3c), but the long term trend indicates drying in the equatorial region and increasing moisture in the southern area (Fig. A3d). Modelled canopy surface moisture in the North America region varied between 0 and 655.1 g $m^{-2}$ (median 0.0, interquartile range 0.0 – 7.2 g $m^{-2}$). Over the entire 60 year period of analysis, canopy moisture was above zero 33.3 % of the time during the spring months (February to May), and 31.6 % of the time during the rest of the year. For East Africa, canopy moisture was above zero 33.6 % of the time during the growing season, and 18.4 % of the time during the rest of the year.

We found strong positive correlations between mean growing season r(*T*) among models, and long-term trends, for both wheat stripe rust in North America and common maize rust in East Africa (Figs. A4-7). Correlations were stronger for the North American region than the East African region (Figs. A6, A7). Mean wheat stripe rust r(*T*) showed a latitudinal trend, declining toward the north (Fig. A4a-c), and increasing over time over most of the region except the far south (Fig. A4d-f). There was no clear latitudinal pattern for common maize rust in East Africa (Fig. A5a-c), but a clear trend of increasing r(*T*) in the south and declining rate in the north over the period of analysis (Fig. A5d-f).

**Discussion**

We found that infection models which consider temperature responses alone closely align with large-scale spatiotemporal patterns produced by models that restricted infection to wet periods. In particular, estimates of long-term mean and trends in r(*T*) calculate at 3-hourly resolution are strongly correlated with those calculated from monthly mean temperatures alone. These results give confidence that our conclusions concerning future shifts in r(*T*), derived from projections of monthly mean temperatures, are robust to potential changes in global humidity patterns. A previous modelling study, on soybean rust, also concluded that humidity and leaf wetness are likely to have negligible effects on disease predictions, because humidity tends to remain high enough for disease development during the crop growing season[11]. Similarly, mean annual temperature is a much stronger determinant of soil-borne fungal plant pathogen abundance than is precipitation[12]. Potato late blight models utilizing monthly microclimate estimates were able to replicate key features of process-based models using hourly microclimate estimates [13]. At the global scale, temperature isoclines are projected to shift latitudinally on land as the climate warms[14], while humidity changes are less certain but unlikely to follow a simple latitudinal trajectory[15]. Hence, modelled changes in infection rate will be dominated by latitudinal shifts in average temperatures.

#### Appendix Figures

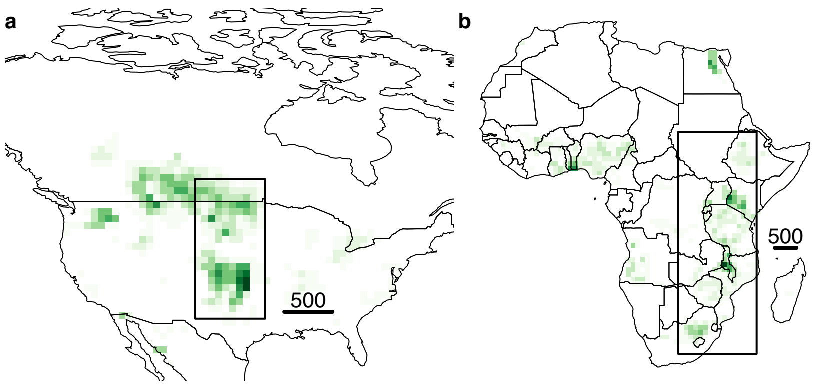

**Fig. A1. Regions for infection model comparison**. (a) North American wheat belt. (b) East Africa maize growing region. Black boxes indicate the modelled region. Green colour density indicates area density of wheat and maize, respectively, in the SPAM 2010 v. 2 global crop allocation model (available from http://mapspam.info) aggregated to 1° resolution. Black scale bar indicates 500km. Thin black lines indicate international borders.

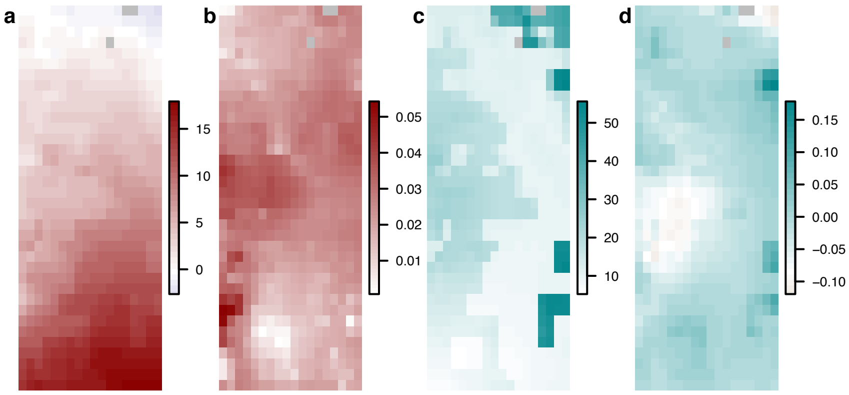

**Fig. A2. Climate summary for North America region, February-May 1958-2017**. (a) Mean temperature (°C). (b) Temperature trend (°C $y^{-1}$). (c) Mean canopy surface moisture (g $m^{-2}$). (d) Canopy surface moisture trend (g $m^{-2}$ $y^{-1}$).

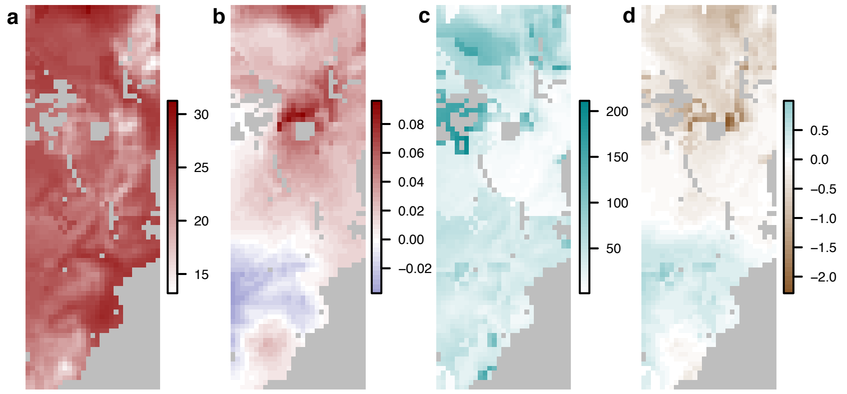

**Fig. A3. Climate summary for East Africa region, February-May 1958-2017**. (a) Mean temperature (°C). (b) Temperature trend (°C $y^{-1}$). (c) Mean canopy surface moisture (g $m^{-2}$). (d) Canopy surface moisture trend (g $m^{-2}$ $y^{-1}$).

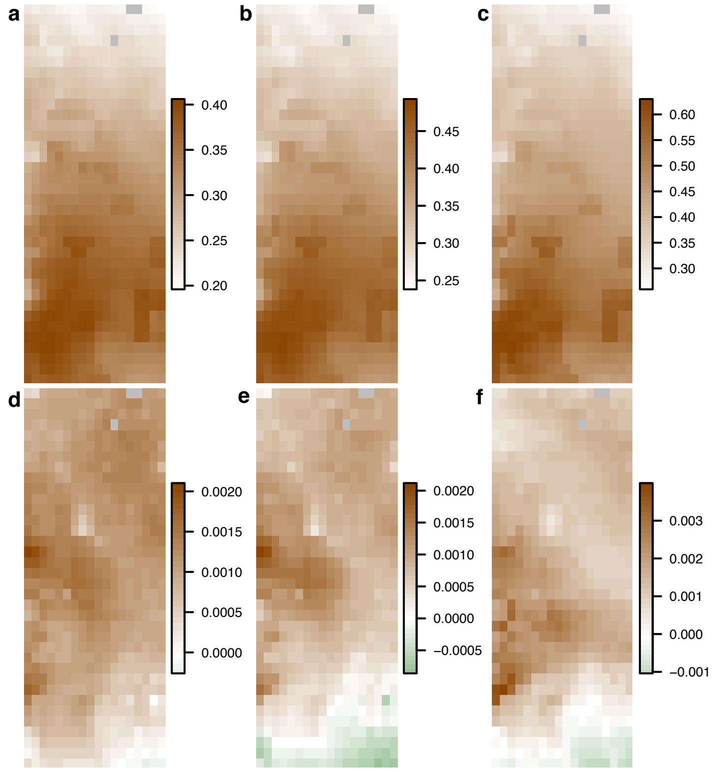

**Fig. A4. North America stripe rust infection rate summary**. Mean rate for (a) wet 3-hourly periods ($r_{wet}$), (b) all 3-hourly periods ($r_{all}$) and (c) mean monthly temperatures ($r_{mon}$). Annual trends of (d) $r_{wet}$ , (e) $r_{all}$ and (f) $r_{mon}$. Infection risk calculated only during spring period (February to May, inclusive).

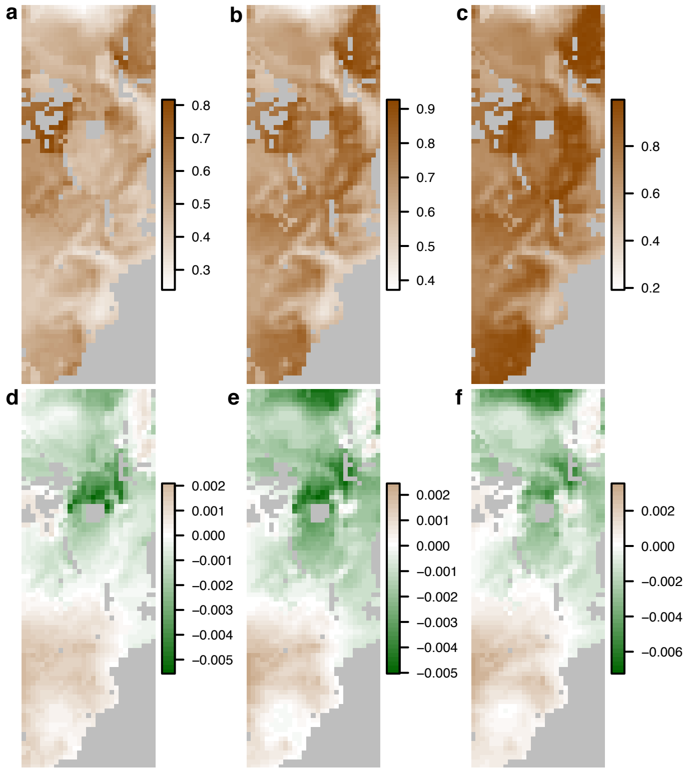

**Fig. A5. East Africa common maize rust infection rate summary**. Mean rate for (a) wet 3-hourly periods ($r_{wet}$ model), (b) all 3-hourly periods ($r_{all}$ model) and (c) mean monthly temperatures ($r_{mon}$ model). Annual trends from (d) $r_{wet}$ model, (e) $r_{all}$ model and (f) $r_{mon}$ model. Infection risk calculated only during growing season obtained from MIRCA2000 model.

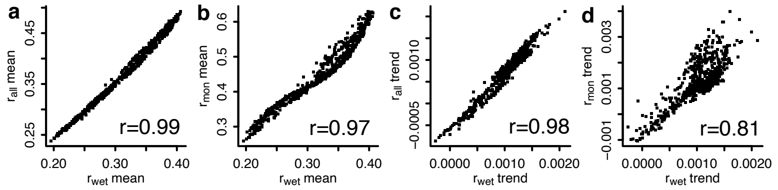

**Fig. A6. Comparison of models for North America region**. a) $r_{all}$ mean vs. $r_{wet}$ mean, (b) $r_{mon}$ mean vs $r_{wet}$ mean, (c) $r_{all}$ annual trend vs. $r_{wet}$ annual trend, (d) $r_{mon}$ trend vs. $r_{wet}$ trend. Points indicate individual grid cells (N=648). Pearson product-moment correlation coefficients are given in the bottom right of each panel.

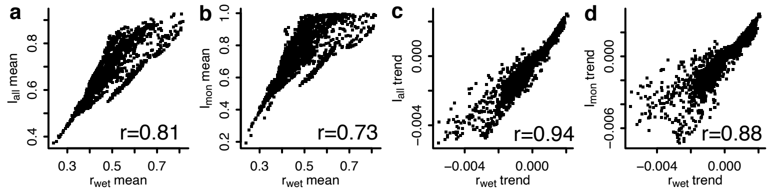

**Fig. A7. Comparison of models for East Africa region**. a) $r_{all}$ mean vs. $r_{wet}$ mean, (b) $r_{mon}$ mean vs $r_{wet}$ mean, (c) $r_{all}$ annual trend vs. $r_{wet}$ annual trend, (d) $r_{mon}$ trend vs. $r_{wet}$ trend. Points indicate individual land surface grid cells (N=1909). Pearson product-moment correlation coefficients are given in the bottom right of each panel.
